# OPA1 Deletion in Brown Adipose Tissue Improves Thermoregulation and Systemic Metabolism via FGF21

**DOI:** 10.1101/2021.01.04.425181

**Authors:** Renata O. Pereira, Alex Marti, Angela C. Olvera, Satya M. Tadinada, Sarah H. Bjorkman, Eric T. Weatherford, Michael Westphal, Pooja H. Patel, Ana K. Kirby, Rana Hewezi, William Bùi Trần, Luis M Garcia Pena, Rhonda A. Souvenir, Monika Mittal, Christopher M. Adams, Matthew J. Potthoff, E. Dale Abel

**Affiliations:** Fraternal Order of Eagles Diabetes Research Center and Division of Endocrinology and Metabolism, Roy J. and Lucille A. Carver College of Medicine, University of Iowa, Iowa City, IA, USA; Department of Obstetrics and Gynecology, Reproductive Endocrinology and Infertility, University of Iowa Hospital and Clinics, Iowa City, IA, USA; Department of Neuroscience and Pharmacology, Roy J. and Lucille A. Carver College of Medicine, University of Iowa, Iowa City, IA, USA

**Keywords:** Brown adipose tissue, thermogenesis, optic atrophy 1, browning, fibroblast growth factor 21, activating transcription factor 4

## Abstract

Adrenergic stimulation of brown adipocytes alters mitochondrial dynamics, including proteolytic processing of the mitochondrial fusion protein optic atrophy 1 (OPA1). However, direct mechanisms linking OPA1 to brown adipose tissue (BAT) physiology are incompletely understood. By deleting OPA1 selectively in BAT (OPA1 BAT KO), we demonstrate that OPA1 is required for cold-induced thermogenesis. Unexpectedly, OPA1 deficiency induced fibroblast growth factor 21 (FGF21) as a BATokine in an activating transcription factor 4 (ATF4)- dependent manner. BAT-derived FGF21 mediates an adaptive response in OPA1 BAT KO mice, by inducing browning of white adipose tissue (WAT), increasing resting metabolic rates, and improving thermoregulation. However, FGF21 does not mediate the resistance to diet-induced obesity observed in these animals. These findings reveal a requirement for OPA1 in BAT thermogenesis, and uncovers a homeostatic mechanism of BAT-mediated metabolic protection governed by an ATF4-FGF21 axis, that is activated independently of BAT thermogenic function.

## Introduction

The prevalence of obesity and its comorbidities, such as type 2 diabetes (T2DM) and cardiovascular disease, has increased dramatically in recent decades (1, 2). Most currently available pharmacological approaches to combat obesity act by reducing caloric intake or impairing fat absorption. However, effective and safe therapies to increase energy expenditure are lacking. The discovery of active brown adipose tissue (BAT) in adult humans has incited interest in exploring BAT activation as a potential strategy for increasing energy expenditure to mitigate obesity and its complications (3-5). Furthermore, BAT has been increasingly recognized as a secretory organ, promoting the release of endocrine factors, or BATokines, that may regulate systemic metabolic homeostasis (6). Therefore, increased understanding of mechanisms regulating BAT thermogenesis and its secretory function could identify new therapeutic strategies for treating obesity and its comorbidities.

Recent studies demonstrated a critical role of mitochondrial dynamics for thermogenic activation of BAT (7, 8). Mitochondrial dynamics describes the process by which mitochondria undergo repeated cycles of fusion and fission. It is mediated by several proteins, including the outer mitochondria membrane fusion proteins mitofusins 1 and 2 (Mfn1 and Mfn2), the inner mitochondrial membrane fusion protein optic atrophy 1 (Opa1), and the fission protein dynamin related protein 1 (Drp1) (9). Norepinephrine treatment induces complete and rapid mitochondrial fragmentation in cultured brown adipocytes through protein kinase A (PKA)-dependent Drp1 phosphorylation, which increases fatty acid oxidation and amplifies thermogenesis (7, 8). Mfn2 is also required for BAT thermogenesis, with its absence rendering mice cold-intolerant, but surprisingly resistant to diet-induced insulin resistance (IR) (10). An earlier study demonstrated that siRNA-mediated knock down of OPA1 in brown adipocytes resulted in a modest, but significant reduction in palmitate oxidation, suggesting a potential role for OPA1 in regulating thermogenesis (11). Moreover, indirect evidence in mice lacking the ATP-independent metalloprotease OMA1, which plays an essential role in the proteolytic inactivation of OPA1, supports the notion that OPA1 regulation of fission is important for thermogenesis. Germline OMA1 deficient mice exhibit increased adiposity, decreased energy expenditure and impaired thermogenesis (11). However, given the ubiquitous expression of OPA1 and OMA1, it is impossible to determine from this model the specific contribution of OPA1 to BAT physiology.

In the present study, we investigated the requirement of OPA1 for BAT’s adaptation to thermogenic stimuli *in vivo*, by generating mice with BAT-specific ablation of the *Opa1* gene (OPA1 BAT KO mice). Our data demonstrated that lack of OPA1 reduced the activation of the thermogenic gene program in BAT, while surprisingly inducing the expression and secretion of fibroblast growth factor 21 (FGF21) as a BATokine, via an ATF4-dependent mechanism. BAT-derived FGF21 mediates an adaptive response characterized by increased browning of white adipose tissue, elevated resting metabolic rates and improved thermoregulation. Nonetheless, FGF21 was dispensable for the resistance to diet-induced obesity (DIO) and insulin resistance (IR) observed in these mice.

## Results

### OPA1 deficiency leads to mitochondrial dysfunction in BAT, while improving energy balance and thermoregulation in mice

To determine the role of OPA1 in BAT physiology, we examined changes in OPA1 levels in BAT in response to high-fat feeding or cold stress. Twelve weeks of high-fat diet (HFD) increased OPA1 mRNA (Figure 1A) and protein levels in BAT (Figure 1B). Three days of cold-exposure (4°C) induced *Opa1* mRNA in BAT (Figure 1C), however, OPA1 total protein levels were significantly reduced (Figure 1D). Inner membrane-anchored long OPA1 (L-OPA1) undergoes proteolytic cleavage by OMA1 resulting in short OPA1 (S-OPA1), that promotes mitochondrial fission (12). Cold temperatures reduced the ratio of L-OPA1 versus the S-OPA1 (Figure 1D), indicating increased proteolytic cleavage of OPA1. Mice deficient for OPA1 specifically in BAT (OPA1 BAT KO) were generated to further investigate the requirement of OPA1 for BAT function. *Opa1* mRNA and protein levels were reduced by 10-fold in BAT (Figure 1E, F), but were maintained in other tissues such as white adipose tissue (WAT), liver and skeletal muscle (Figure S1A-C). OPA1-deficient BAT revealed increased numbers of enlarged unilocular lipid droplets, suggesting whitening of BAT (Figure 1G). Ultrastructurally, mitochondria appeared more fragmented and lamellar cristae structure was disrupted (Figure 1G). Isolated mitochondria from OPA1-deficient BAT revealed impaired mitochondrial bioenergetics, exemplified by reduced pyruvate-malate (Figure 1H), and palmitoyl-carnitine dependent oxygen consumption (Figure 1I) and ATP synthesis rates (Figure 1J). Mitochondrial dysfunction in BAT has been associated with reduced metabolic rates and increased fat accumulation in mice (13, 14). Unexpectedly, OPA1 BAT KO mice exhibited a small, but significant reduction in body mass, which became more striking with age (Figure 1K). Total fat mass was unchanged and total lean mass was reduced in 8-week-old KO mice (Figure 1L), whereas percent fat mass and lean mass relative to body weight were not significantly changed (Figure S1D and E). However, the expected age-dependent increase in total fat mass and lean mass was strikingly attenuated in 20-week-old KO mice, which had reduced total fat mass and lean mass, relative to WT mice (Figure 1M). Percent fat mass was reduced, while percent lean mass was increased in 20-week-old KO mice (Figure S1F and G). Although fat mass was unchanged in 8-week-old KO mice, BAT mass was significantly increased at this age, but significantly reduced by 20-weeks (Figure 1N). In contrast, weight of inguinal (Figure 1O) and gonadal (Figure 1P) white adipose tissue depots (gWAT and iWAT, respectively) were significantly reduced in KO mice at both 8 and 20 weeks of age. Despite these changes in body composition, glucose tolerance was unaffected in 8-week (Figure S1H and I) and 20-week-old KO mice (Figure S1J and K). The reduction in body weight and fat mass persisted in aging mice, as 50-week-old female mice exhibited an approximate 40% reduction in body mass (Figure S1L), a 3-fold reduction in body fat (Figure S1M) and a significant increase in lean mass (Figure S1N), relative to body weight. At this age, glucose tolerance was significantly improved in KO mice relative to WT mice (Figure S1O and P). To further elucidate mechanisms responsible for the weight loss, 8-week old mice were studied in metabolic chambers. At thermoneutrality (30°C), oxygen consumption was significantly increased in KO mice during both light and dark cycles (Figure 1Q), which likely contributes to the lean phenotype in KO mice, as no changes were detected in food intake (Figure S1Q) or locomotor activity (Figure S1R) between genotypes. Despite reduced mitochondrial fatty acid oxidation in BAT, core body temperature was significantly increased in KO mice (Figure 1R), indicating increased thermogenesis, even at 30°C. To determine if these phenotypic changes would persist in the absence of lifelong thermogenic activation, a separate cohort of mice was raised at 30°C. Body mass remained reduced in KO mice compared to WT mice (Figure S2A), while percent fat mass (Figure S2B) and lean mass (Figure S2C) were unchanged between genotypes. As observed in mice raised at room temperature (22°C), BAT mass (Figure S2D) was increased, whereas gWAT (Figure S2E) and iWAT mass (Figure S2F) were significantly reduced in KO mice reared at 30°C. Resting metabolic rates remained elevated in KO mice raised at 30°C, and were significantly increased during the light cycle (Figure S2G). Food consumption was unchanged between WT and KO mice (Figure S2H), but locomotor activity (Figure S2I) was significantly reduced in KO mice during the dark cycle. Together, our data strongly suggest that the improved energy balance and thermoregulation observed in OPA1 BAT KO mice occur independently of BAT thermogenic activation.

**Figure 1:**
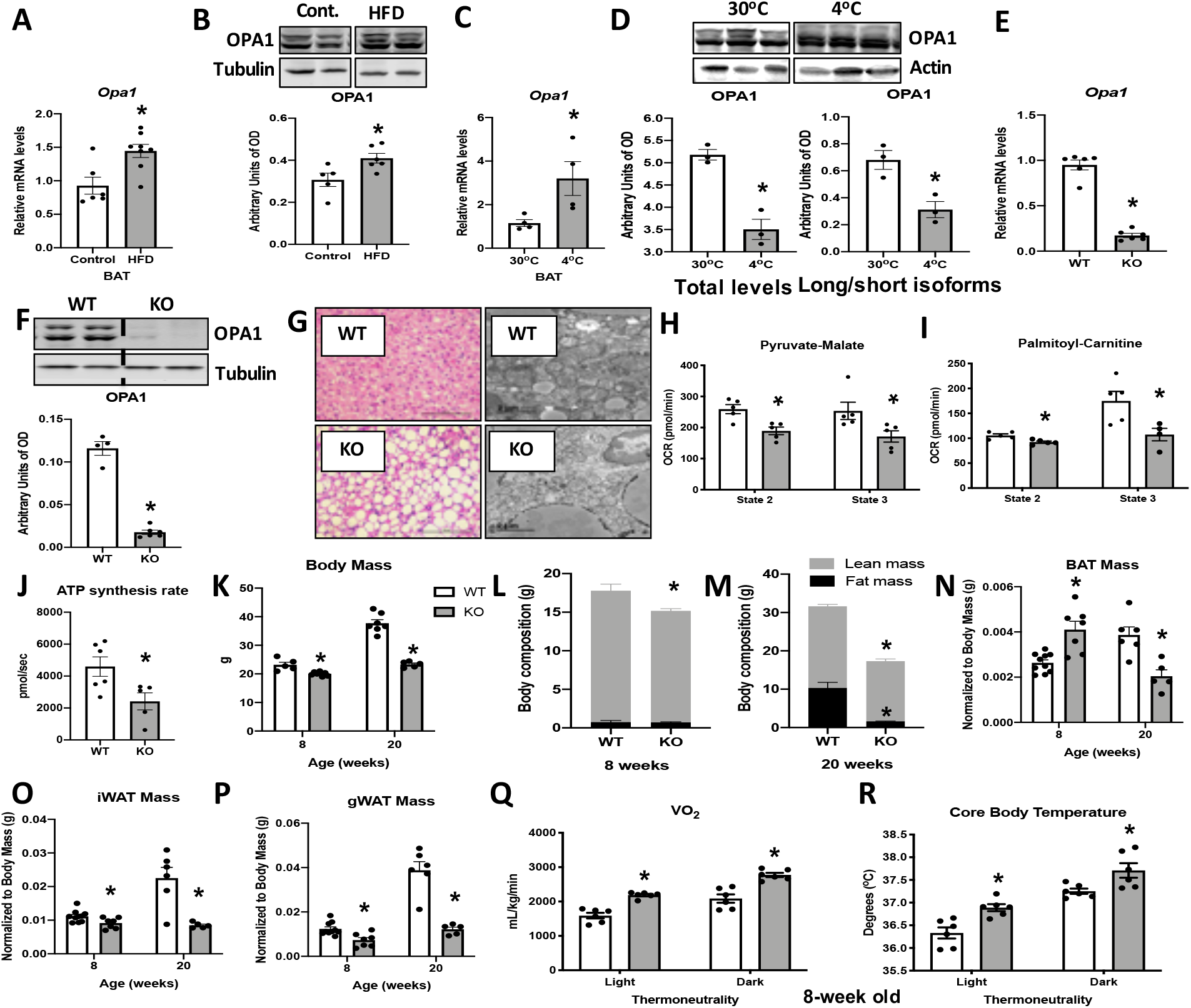
OPA1 deficiency leads to mitochondrial dysfunction in BAT, while improving energy balance and thermoregulation in mice. **A-B**. OPA1 expression in brown adipose tissue (BAT) of wild type (WT) mice fed either control (10% fat) or a high-fat diet (HFD 60% fat) for 12 weeks. **A**. *Opa1* mRNA expression in BAT. **B**. Representative immunoblot of OPA1 and densitometric analysis of OPA1 normalized by tubulin (images were cropped from the same membrane). **C-D**. OPA1 expression in BAT of WT mice maintained at 30°C or 4°C for 3 days. **C**. *Opa1* mRNA expression in BAT. **D**. Representative immunoblot of OPA1 and densitometric analyses of OPA1 (total levels and long/short isoforms - images were cropped from the same membrane). **E-J**. Morphological and functional characterization of 8-week-old OPA1 BAT KO mice (KO). **E**. *Opa1* mRNA expression in BAT. **F**. Representative immunoblot of OPA1 and densitometric analysis of OPA1 normalized to tubulin (dashed line separates genotypes). **G**. Representative images of H&E-stained histological sections and electron micrographs of BAT from 8-week-old WT and KO mice (n=3). Scale bar = 100μm and 2μm, respectively. **H-I**. Functional analysis of mitochondria isolated from BAT of WT and KO mice. **H**. Basal (state 2) and ADP-stimulated (state 3) pyruvate-malate-supported oxygen consumption rates (OCR). **I**. State 2 and state 3 palmitoyl-carnitine-supported OCR. **J**. Palmitoyl-carnitine-supported ATP synthesis rates. **K-P**. Body mass and body composition in 8 and 20-week old WT and KO mice. **K**. Body mass (8 and 20 weeks of age). **L**. Body composition (8 weeks of age). **M**. Body composition (20-weeks of age). **N**. BAT mass. **O**. Inguinal white adipose tissue (iWAT) mass. **P**. Gonadal white adipose tissue mass (gWAT). **Q**. Oxygen consumption in 8-week-old mice housed at 30°C. **R**. Core body temperature in 8-week-old mice housed at 30°C. Data are expressed as means ± SEM. Significant differences were determined by Student’s *t*-test, using a significance level of *P* < 0.05. (*) Significantly different vs. WT mice.

### OPA1 BAT KO mice exhibit improved tolerance to cold despite impaired thermogenic activation of BAT

Impaired mitochondrial function in BAT has been linked to cold-intolerance in mice (10, 11, 14). To determine the response of OPA1 BAT KO to cold stress, we first measured rectal body temperature during acute cold exposure. Body temperature declined at a slower rate in KO mice relative to WT mice during 4 hours of cold exposure (Figure 2A). Next, we prolonged the thermal challenge by exposing a separate cohort of mice to 4°C for 3 days. mRNA analysis revealed increased expression of the thermogenic genes *Ucp1* (Figure 2B), *Prdm16* (Figure 2C), and *Ppargc-1α* (Figure 2D) in BAT of WT mice after 3 days of cold exposure, which was significantly attenuated in KO mice. Nonetheless, core body temperature was elevated in KO mice, and was significantly increased in the dark cycle (Figure 2E). Thus, mice lacking OPA1 in BAT have improved thermoregulatory capacity despite BAT dysfunction. Whole animal oxygen consumption remained higher in KO mice after 3 days of cold exposure (Figure 2F), despite the absence of any increase in locomotor activity (Figure 2G) or food intake during the dark cycle (Figure 2H), which contrast with responses of WT mice.

**Figure 2:**
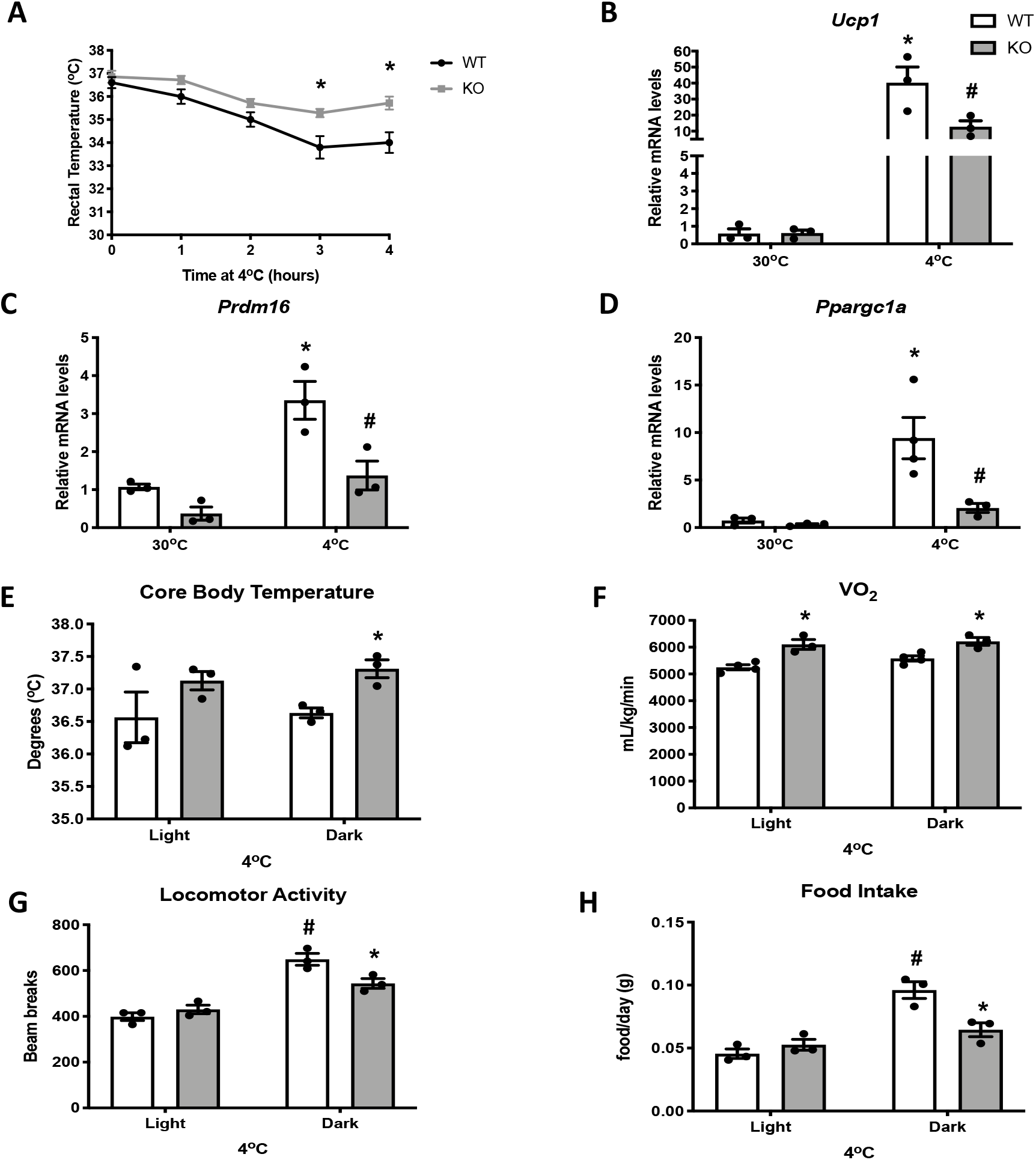
OPA1 BAT KO mice exhibit improved tolerance to cold despite impaired thermogenic activation of BAT. **A**. Rectal temperature in 8-week-old WT and KO mice exposed to acute cold stress (4°C) over the period of 4 hours. **B-D**. mRNA expression of thermogenic genes in BAT of WT and KO mice housed at 30°C or 4°C for 3 days. **B**. Relative *Ucp1* mRNA levels. **C**. Relative *Prdm16* mRNA levels. **D**. Relative *Ppargc1α* mRNA levels. mRNA expression was normalized to *Gapdh*. **E-F**. Indirect calorimetry and core body temperature in WT and KO mice exposed to 4°C for 3 days. **E**. Core body temperature. **F**. Oxygen consumption. **G**. Locomotor activity. **H**. Food intake (average for each cycle). Data are expressed as means ± SEM. Significant differences were determined by Two-Way ANOVA using a significance level of *P* < 0.05. (*) Significantly different vs. WT mice or vs. 30°C, (#) significantly different from light cycle or WT mice at 4°C.

### OPA1 deletion in BAT results in compensatory browning of WAT

To determine the presence of compensatory browning of WAT in OPA1 BAT KO mice, we performed morphological and biochemical analysis of the inguinal fat depots (iWAT), which are prone to undergoing browning in mice. Histologically, iWAT morphology in KO mice revealed several regions presenting smaller adipocytes with multilocular lipid droplets that resembled brown adipocytes (Figure 3A). Immunohistochemistry (Figure 3A) and immunoblots of mitochondrial protein (Figure 3B) both revealed significant induction of uncoupling protein 1 (UCP1) in KO mice. Surprisingly, OPA1 protein levels were also increased in mitochondria isolated from iWAT (Figure 3B). Because the specific UCP1 Cre mouse used in the present study has been shown to promote recombination of floxed alleles in both brown and beige adipocytes, particularly in response to cold (15, 16), we examined *Cre* expression in iWAT of WT and KO mice, relative to BAT. Cre expression in the BAT of KO mice was ∼ 300-fold higher than its expression in iWAT, which was not statistically increased from WT iWAT (Figure S3A). These data suggest that *Cre* expression in iWAT is insufficient to promote recombination and deletion of OPA1 at ambient temperature conditions in this model. Mitochondria ultrastructure in beige iWAT was characterized by abundant and tightly organized lamellar-cristae in KO mice relative to WT mice (Figure 3C). Transcriptionally, mRNA expression of thermogenic and fatty acid oxidation genes (Figure 3D) was induced in iWAT of KO mice, which correlated with increased pyruvate-malate (Figure 3E) and palmitoyl-carnitine supported oxygen consumption rates (Figure 3F) in isolated mitochondria. As reported in other models of BAT dysfunction (17, 18), we observed increased sympathetic activity in iWAT of KO mice, as measured by increased protein levels of tyrosine hydroxylase relative to WT mice (Figure 3G).

**Figure 3:**
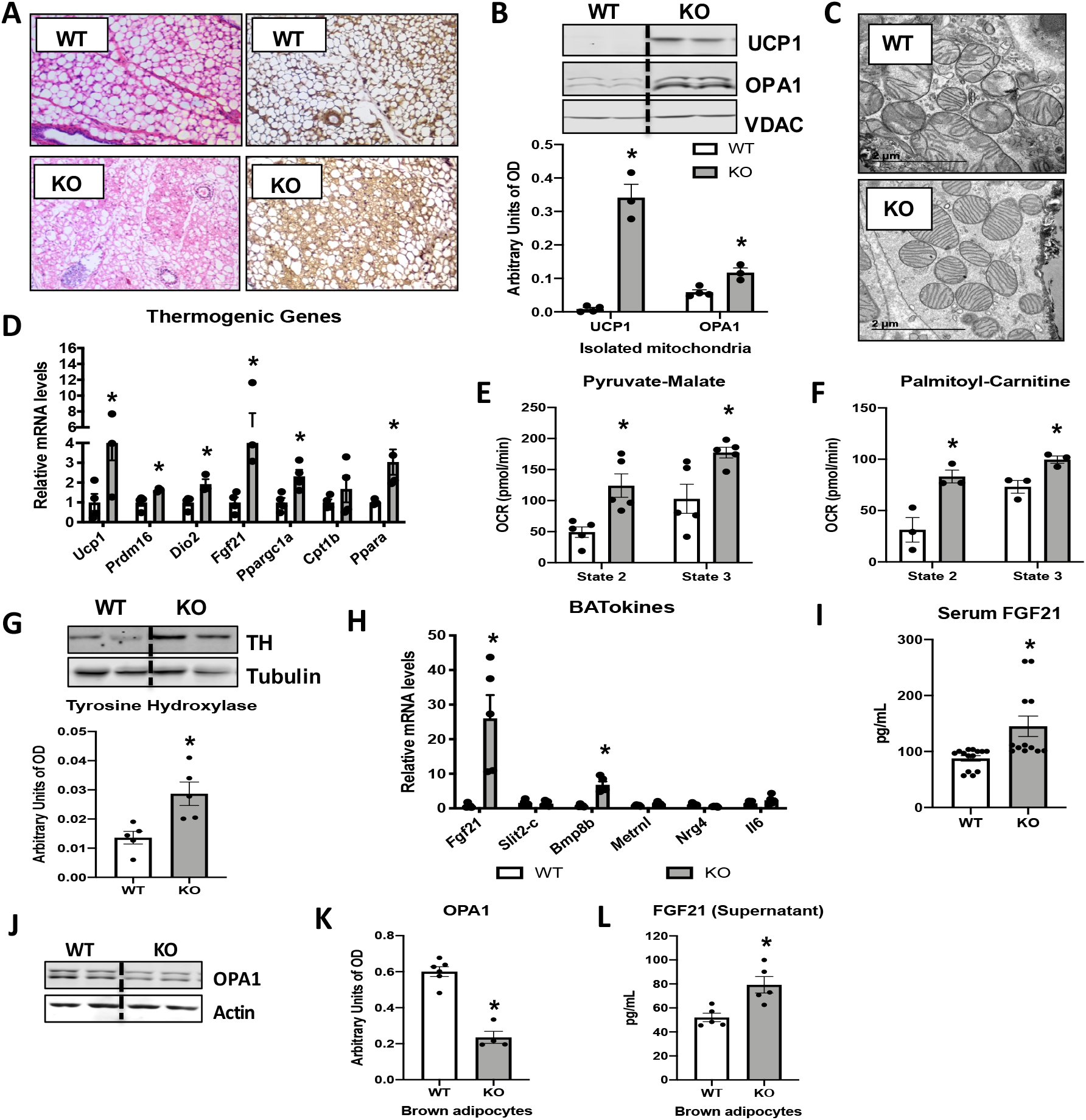
OPA1 deletion in BAT results in compensatory browning of WAT. **A-G:** Morphological and functional characterization of inguinal white adipose tissue (iWAT) in 8-week old WT and KO mice. **A**. Representative iWAT sections stained with H&E or after immunohistochemistry against UCP1. Scale bar = 100μm (n=3). **B**. Representative immunoblot (dashed line separates genotypes) and densitometric analysis of UCP1 and OPA1 in mitochondria isolated from iWAT normalized to VDAC. **C**. Representative electron micrographs of iWAT from WT and KO mice. Scale bar = 2 µm (n=3). **D**. mRNA expression of thermogenic genes. **E-F**. Functional analysis of mitochondria isolated from iWAT. **E**. State 2 and state 3 pyruvate-malate-supported mitochondrial OCR. **F**. State 2 and state 3 palmitoyl-carnitine-supported mitochondrial OCR. **G**. Representative immunoblot (dashed line separates genotypes) and densitometric analysis of tyrosine hydroxylase (TH) normalized to tubulin. **H**. mRNA levels of BATokines in BAT extracts from 8-week-old WT and KO mice. **I**. Serum levels of fibroblast growth factor 21 (FGF21) in 8-week old WT and KO mice. **J**. Representative immunoblots of OPA1 normalized to actin in primary brown adipocytes (dashed line separates genotypes). **K**. Densitometric analysis of OPA1 normalized to tubulin in brown adipocytes. **L**. FGF21 levels measured in the culture media collected from WT and OPA1-deficient brown adipocytes. Data are expressed as means ± SEM. Significant differences were determined by Student’s *t*-test, using a significance level of *P* < 0.05. (*) Significantly different vs. WT mice.

To investigate the role of brown adipokines or ‘BATokines’ in the adaptations observed in OPA1 BAT KO mice, we investigated the mRNA expression of a sub-set of previously described BATokines. Of these targets, fibroblast growth factor 21 (FGF21) was highly induced in BAT of KO mice (Figure 3H), which correlated with increased circulating concentrations of FGF21 in KO mice under ad libitum-fed conditions (Figure 3I) or after a 6-hour fast (Figure S3B). Notably, *Fgf21* mRNA expression was reduced in the liver (Figure S3C), suggesting that BAT, rather than liver, contributed to FGF21 circulating levels in KO mice. Short-term knockdown of OPA1 in brown adipocytes (Figure 3J and K) was sufficient to increase the release of FGF21 (Figure 3L) into the cell culture media, demonstrating that OPA1 deletion induces FGF21 secretion in a cell-autonomous manner. Thus, OPA1 deficiency in brown adipocytes induces secretion of FGF21 independently of thermogenic activation.

### BAT-derived FGF21 is required for increased resting metabolic rates and improved thermoregulation in mice lacking OPA1 in BAT during isocaloric feeding

To determine if BAT-derived FGF21 mediates the systemic metabolic adaptations in OPA1 BAT KO mice, we generated mice lacking both OPA1 and FGF21 specifically in BAT (DKO mice). mRNA expression of *Opa1* and *Fgf21* was significantly reduced in BAT of DKO mice (Figure 4A), which completely normalized circulating FGF21 levels (Figure 4B). Pyruvate-malate (Figure S4A) and plamitoyl-carnitine (Figure S4B) supported state 2 and state 3 mitochondrial respirations were reduced in the BAT of DKO mice to the same extent as observed in OPA1 BAT KO mice. Furthermore, mRNA expression of thermogenic genes was repressed in the BAT of DKO mice (Figure S4C). Body mass was normalized in DKO mice (Figure 4C), and fat mass and lean mass were unchanged (Figure 4D and E) between 8-week-old DKO mice and age-matched WT controls. Similar to OPA1 BAT KO mice, BAT mass was increased (Figure 4F) in DKO mice at 8-week of age, reiterating that FGF21 does not impact the BAT phenotype in this model. In contrast, the reduction in gWAT and iWAT mass observed in OPA1 BAT KO mice was no longer detectable in DKO mice (Figure 4G and H). There were no statistically significant changes in oxygen consumption between DKO mice and their WT littermate controls (Figure 4I).

**Figure 4:**
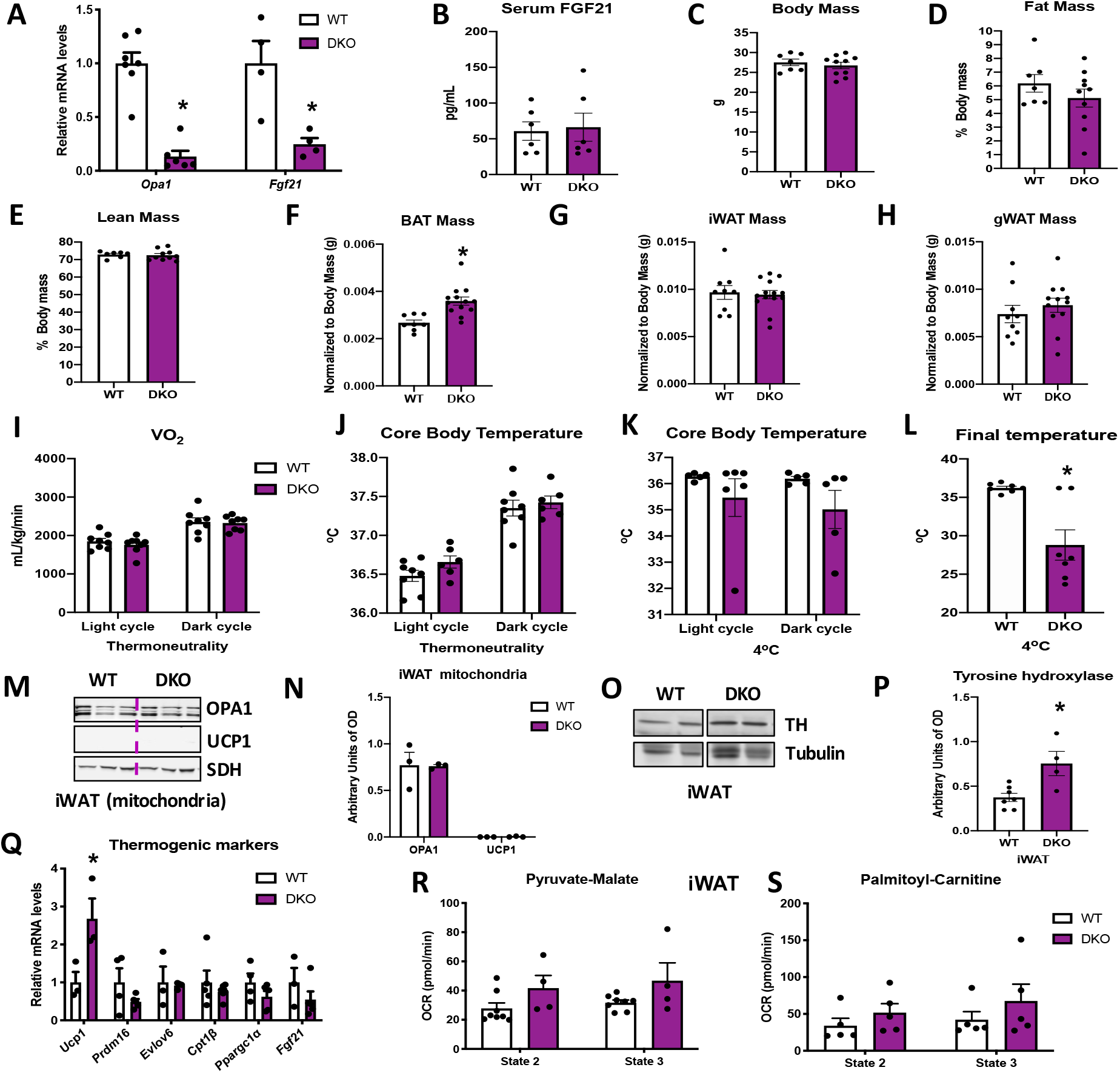
BAT-derived FGF21 is required for increased resting metabolic rates and improved thermoregulation in mice lacking OPA1 in BAT during isocaloric feeding. **A-L**. Data characterizing 8-12-week-old OPA1/FGF21 DKO mice. **A**. mRNA expression of *Opa1* and *Fgf21* in BAT of DKO mice. **B**. FGF21 serum levels collected under ad libitum-fed conditions. **C**. Total body mass. **D**. Percent fat mass. **E**. Percent lean mass. **F**. BAT mass. **G**. iWAT mass. **H**. gWAT mass. **I**. Oxygen consumption (30°C). **J**. Core body (30°C). **K**. Core body temperature (4°C) (data is represented as average core body temperature during the light and dark cycles over three days of continuous monitoring). **L**. Final core body temperature recorded for each individual mouse (4°C). **M-R**. Data of iWAT from 8-week old DKO mice. **M**. Representative immunoblot for OPA1 and UCP1 in isolated mitochondria (dashed line separates genotypes). **N**. Densitometric analysis of OPA1 and UCP1 protein levels normalized to succinate dehydrogenase (SDH). **O**. Representative immunoblot for tyrosine hydroxylase (TH) in iWAT (images were cropped from the same membrane). **P**. Densitometric analysis of TH protein levels normalized to tubulin. **Q**. mRNA expression of thermogenic genes in iWAT. **R-S**. Functional analysis of mitochondria isolated from iWAT. **R**. State 2 and state 3 pyruvate-malate-supported mitochondrial OCR. **S**. State 2 and state 3 palmitoyl-carnitine-supported mitochondrial OCR. Data are expressed as means ± SEM. Significant differences were determined by Student’s *t*-test or Two-Way ANOVA, using a significance level of *P* < 0.05. (*) Significantly different vs. WT mice.

Locomotor activity (Figure S4D) and food consumption (Figure S4E) were also unchanged between genotypes at 30°C.

The increase in baseline core body temperature observed in OPA1 BAT KO mice, was completely lost in DKO mice, suggesting that BAT-derived FGF21 mediates the increase in core body temperature at 30°C (Figure 4J). Moreover, oxygen consumption was not increased in DKO mice during cold exposure (Figure S4F). Core body temperature tended to be lower after 3 days of cold exposure, but did not reach statistical significance (Figure 4K). However, the last temperature recorded by telemetry for each mouse was significantly reduced in DKO mice, and better reflects the defect in thermoregulation observed in these mice (Figure 4L). This was primarily due to a steep reduction in core body temperature in 4 out of 7 DKO mice, which died of hypothermia before the end of the cold exposure studies (data not shown). Locomotor activity was unchanged between genotypes (Figure S4G), while food intake was significantly reduced in DKO mice during the dark cycle, likely due to hypothermia (Figure S4H). FGF21 has been implicated in browning of WAT (19). We, therefore, measured browning markers in iWAT of DKO mice. UCP1 and OPA1 protein levels (Figure 4M) were unchanged in mitochondria isolated from iWAT of DKO mice (Figure 4M and N). Of note, although tyrosine hydroxylase protein levels remained elevated in iWAT of DKO mice (Figure 4O and P), the induction of thermogenic genes in iWAT observed in OPA1 BAT KO mice was absent in DKO mice (Figure 4Q). Furthermore, pyruvate-malate- (Figure 4R) and palmitoyl-carnitine-supported (Figure 4S) state 2 and state 3 mitochondrial respirations were unchanged between mitochondria isolated from iWAT of WT and DKO mice. Thus, BAT-derived FGF21 is required for the compensatory browning and for thermoregulation in OPA1 BAT KO mice fed isocaloric diet.

### OPA1 deletion in BAT prevents diet-induced obesity and insulin resistance

Although impaired BAT thermogenesis is frequently associated with increased susceptibility to diet-induced obesity (DIO) (20-22), we hypothesized that the increased resting metabolic rates could protect OPA1 BAT KO mice from DIO. Indeed, KO mice fed HFD for 12 weeks weighed the same as WT mice fed low-fat control diets (Figure 5A). KO mice fed the control diet weighed significantly less than WT mice fed the same diet (Figure 5A). The reduction in body mass occurred at the expense of fat mass, as percent fat mass (Figure 5B) was significantly reduced and percent lean mass (Figure 5C) was significantly increased in KO mice fed HFD. BAT mass (Figure 5D), gWAT mass (Figure 5E) and iWAT mass (Figure 5F) were also significantly reduced in HFD-fed KO mice. Indirect calorimetry revealed increased oxygen consumption in KO mice compared to WT mice, regardless of the diet (Figure 5G), while food consumption (Figure 5H) and locomotor activity (Figure 5I) were unchanged between genotypes. Thus, increased metabolic rates likely contribute to leanness in these mice. Glucose homeostasis and insulin sensitivity were also improved in KO mice fed HFD relative to WT mice. As expected, glucose tolerance was impaired in HFD-fed WT mice, which was significantly attenuated in KO mice (Figure 5J and K). Fasting glucose levels were increased in WT mice, but not in KO mice fed HFD (Figure 5L). Similarly, insulin tolerance tests (ITT) revealed impaired insulin sensitivity in WT mice fed HFD, that was prevented in KO mice (Figure 5 M and N). Fasting insulin levels, were also significantly reduced in KO mice versus WT mice fed HFD (Figure 5O). Hepatic steatosis was attenuated in KO mice (Figure S5A, B) relative to WT mice fed HFD, and serum triglycerides levels were completely normalized (Figure S5C). Diet-induced thermogenic activation of BAT was significantly attenuated in KO mice, as evidenced by reduced *Ucp1* mRNA levels (Figure S5D); however, *Ucp1* transcript levels were significantly elevated in the iWAT of KO mice upon high-fat feeding, indicating increased browning of iWAT (Figure S5E). Of note, tyrosine hydroxylase levels were significantly reduced in iWAT of KO mice fed HFD, relative to WT mice (Figure S5F).

**Figure 5:**
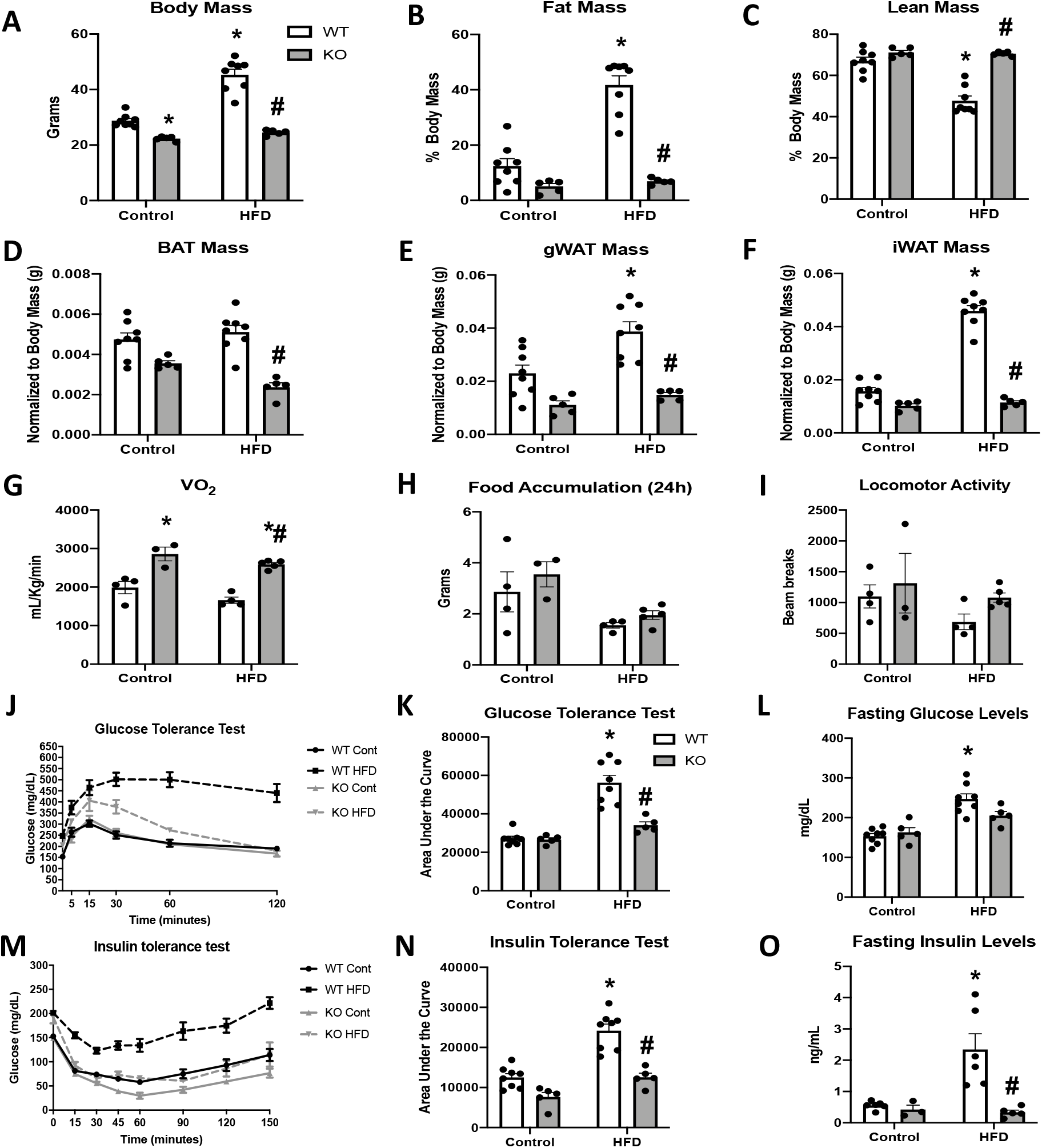
OPA1 deletion in BAT prevents diet-induced obesity and insulin resistance. **A-O**. Data from WT and OPA1 BAT KO mice fed either a control diet (Cont) or a high-fat diet (HFD) for 12 weeks. **A**. Total body mass. **B**. Percent ratio of fat mass to body mass. **C**. Percent ratio of lean mass to body mass. **D**. BAT mass. **E**. gWAT mass. **F**. iWAT mass. **G**. Oxygen consumption. **H**. Food accumulation during a 24-hour period. **I**. Locomotor activity. **J**. Glucose tolerance test (GTT). **K**. Area under the curve for the GTT. **L**. Fasting glucose levels. **M**. Insulin tolerance test (ITT). **N**. Area under the curve for the ITT. **O**. Fasting insulin levels. Data are expressed as means ± SEM. Significant differences were determined by Two-Way ANOVA, using a significance level of *P* < 0.05. (*) Significantly different vs. WT control, (#) significantly different vs. WT HFD.

### BAT-derived FGF21 does not mediate resistance to diet-induced obesity in OPA1 BAT KO mice

To determine if the resistance to DIO required BAT-derived FGF21, we fed OPA1/FGF21 BAT DKO mice either a control or a HFD for 12 weeks. Under control diet conditions, DKO mice lacked the reduction in body mass noted in OPA1 KO mice (Figure 6A). However, when fed a HFD, the increase in total body mass (Figure 6A) and fat mass (Figure 6B) observed in WT mice on a HFD was completely prevented in DKO mice. Furthermore, the percent lean mass relative to body mass was reduced in WT mice fed HFD (Figure 6C). BAT mass was reduced in DKO mice (Figure S6A), and the diet-induced increase in gonadal and inguinal WAT mass (Figure S6B and C) was completely prevented in DKO mice relative to WT controls. Indirect calorimetry confirmed that, in mice fed control diet, oxygen consumption was unchanged in DKO mice relative to WT controls (Figure 6D); however, DKO mice fed a HFD had increased oxygen consumption relative to WT mice (Figure 6D). This increase in metabolic rates likely contributed to their resistance to weight gain, as we observed no significant changes in food intake (Figure 6E) or locomotor activity (Figure 6F) between genotypes, regardless of diet.

**Figure 6:**
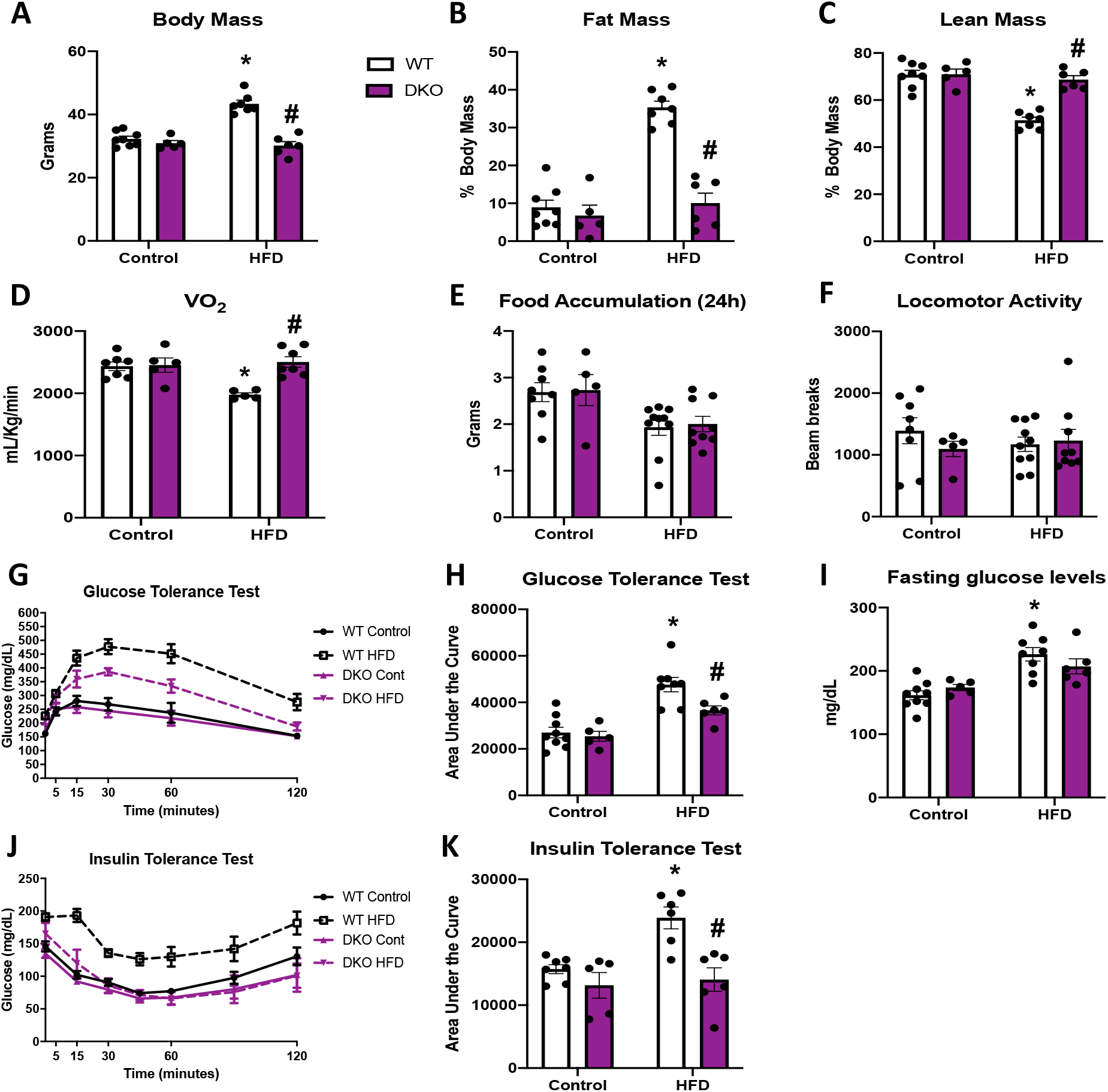
BAT-derived FGF21 does not mediate resistance to diet-induced obesity in OPA1 BAT KO mice. **A-K**. Data on WT and OPA1/FGF21 DKO mice fed either a control diet (Cont) or a high-fat diet (HFD) for 12 weeks. **A**. Total body mass. **B**. Percent ratio of fat mass to body mass. **C**. Percent ratio of lean mass to body mass. **D**. Oxygen consumption. **E**. Food accumulation during a 24-hour period. **F**. Locomotor activity. **G**. Glucose tolerance test. **H**. Area under the curve for the glucose tolerance test. **I**. Fasting glucose levels. **J**. Insulin tolerance test. **K**. Area under the curve for the insulin tolerance test. Data are expressed as means ± SEM. Significant differences were determined by Two-Way ANOVA, using a significance level of *P* < 0.05. (*) Significantly different vs. WT control, (#) significantly different vs. WT HFD.

Furthermore, liver triglycerides levels were significantly reduced in DKO mice relative to WT mice fed a HFD (Figure S6D), consistent with attenuation of diet-induced hepatic steatosis. BAT-derived FGF21 is also dispensable for the improvements in glucose homeostasis and insulin sensitivity in OPA1 BAT KO mice, as HFD-induced glucose intolerance was attenuated in DKO mice (Figure 6G and H), and fasting glucose levels were unchanged (Figure 6I). Finally, insulin sensitivity was also ameliorated in DKO mice compared to WT fed HFD, as shown by the insulin tolerance test (Figure 6J and K). *Ucp1* mRNA levels were significantly increased in BAT (Figure S6E) and iWAT (Supplemental Figure 6F) of DKO mice fed a HFD, relative to mice fed control diet, consistent with diet-induced thermogenic activation and browning. However, tyrosine hydroxylase protein levels were downregulated in iWAT of DKO mice fed HFD, relative the WT mice (Figure S6G). High-fat feeding significantly increased FGF21 serum levels in WT mice, which was prevented in DKO mice (Figure S6H). Although FGF21 circulating levels were slightly elevated in OPA1 BAT KO mice fed a control diet relative to WT mice, the diet-induced increase in FGF21 levels was blunted in KO mice (Figure S6I). Thus, FGF21-independent mechanisms mediate the resistance to DIO and insulin resistance observed in OPA1 BAT KO mice.

### ATF4 is required for FGF21 induction in OPA1 BAT KO mice

OPA1 deletion in BAT induced endoplasmic reticulum (ER) stress, as demonstrated by increased phosphorylation of the Eukaryotic Translation Initiation Factor 2A (eIF2*α*), which promotes selective translation of the activating transcription factor 4 (ATF4) (Figure 7A). Moreover, mRNA expression of the ER stress genes *Atf4, Chop* and *Ire1α* was increased in the BAT of KO mice (Figure 7B). ATF4 binds to the FGF21 promoter to induce its expression in multiple cell types (23-25). We, therefore, generated mice with concurrent deletion of OPA1 and ATF4 in BAT (OPA1/ATF4 BAT DKO) to determine if ATF4 is required for FGF21 induction in KO mice (Figure 7C and D). Indeed, ATF4 deletion in OPA1 BAT KO mice completely normalized FGF21 mRNA (Figure 7E) and circulating levels (Figure 7F). We, then, investigated parameters believed to be regulated by BAT-derived FGF21 in these OPA1/ATF4 BAT DKO mice. Body mass was unchanged in mice lacking ATF4, relative to their WT littermate controls (Figure 7G). BAT mass remained elevated in DKO mice (Figure 7H), while gonadal (Figure 7I) and inguinal (Figure 7J) WAT mass were unchanged between genotypes. DKO mice lacked the increase in resting metabolic rates (Figure 7K) or core body temperature (Figure 7L) observed in OPA1 BAT KO mice at 30°C. Furthermore, activation of thermogenic genes in iWAT was ablated in DKO mice (Figure 7M), indicating that the ATF4-FGF21 axis is required for the baseline compensatory browning of iWAT in OPA1 BAT KO mice. At 4°C, core body temperature tended to be lower in DKO mice, but did not reach statistical significance (Figure 7N). Nonetheless, the last temperature recorded by telemetry for each mouse was significantly reduced in DKO mice (Figure 7O). Similar to observations in OPA1/FGF21 DKO mice, a subset of mice died of cold-induced hypothermia, adding variability to the averaged data. Cold-induced thermogenic activation of BAT (Figure 7P), and activation of browning were attenuated in DKO mice, as demonstrated by reduced mRNA expression of thermogenic genes (Figure 7Q) and reduced UCP1 protein levels (Figure 7R). Thus, concomitant deletion of OPA1 and ATF4 in BAT phenocopies the effects of FGF21 deletion in OPA1 BAT KO mice.

**Figure 7:**
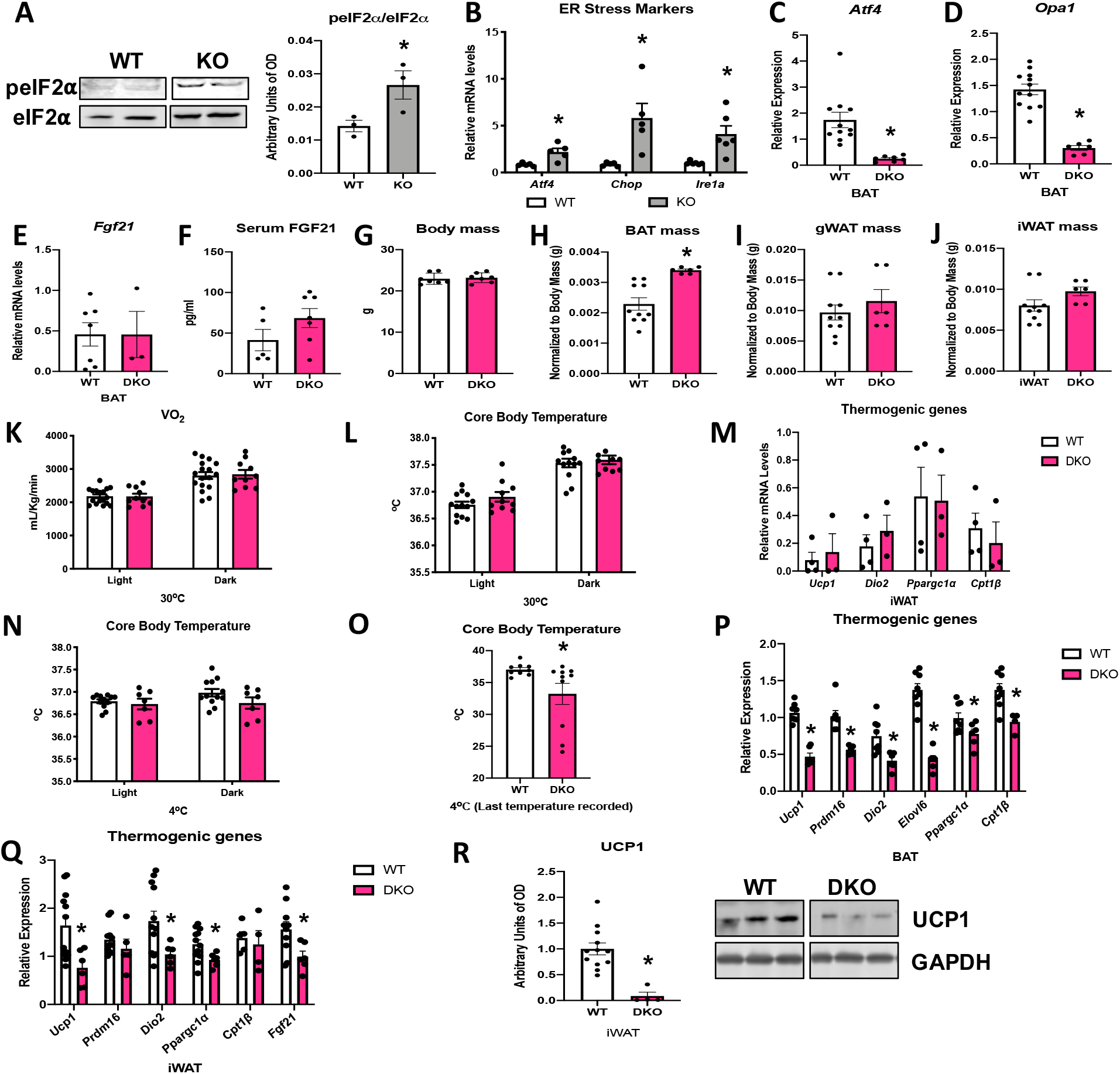
ATF4 is required for FGF21 induction in OPA1 BAT KO mice. **A-B:** Analysis of ER stress in BAT tissue from WT and OPA1 BAT KO mice (KO). **A**. Representative immunoblot for phosphorylated eIF2*α* over total eIF2*α* and respective densitometric quantification (images were cropped from the same membrane). **B**. mRNA expression of ER stress markers. **C-R**. Data collected in 8-10-week old OPA1/ATF4 BAT DKO mice. **C**. mRNA expression of *Atf4* in BAT. **D**. mRNA expression of *Opa1* in BAT. **E**. FGF21 mRNA expression in BAT. **F**. FGF21 serum levels at ambient temperature and ad libitum-fed conditions. **G**. Body mass. **H**. BAT mass normalized to body mass. **I**. gWAT mass normalized to body mass. **J**. iWAT mass normalized to body mass. **K**. Oxygen consumption measured at 30°C. **L**. Core body temperature measured at 30°C. **M**. mRNA expression of thermogenic genes in iWAT samples collected at ambient temperature. **N**. Core body temperature in DKO mice exposed to 4°C (data is represented as average core body temperature during the light and dark cycles over three days of continuous monitoring). **O**. Final core body temperature recorded for each individual mouse (4°C). **P**. mRNA expression of thermogenic genes in BAT samples. **Q**. mRNA expression of thermogenic genes in iWAT samples. **R**. Representative immunoblot for UCP1 normalized to GAPDH (images were cropped from the same membrane) in iWAT and respective densitometric quantification (**P-R** collected after 3 days at 4°C). Data are expressed as means ± SEM. Significant differences were determined by Student’s T-test, using a significance level of *P* < 0.05. (*) Significantly different vs. WT.

## Discussion

Mitochondrial fission contributes to BAT thermogenesis (7, 8). However, the role of OPA1 in BAT function was not well-understood. We provide direct evidence that OPA1 maintains mitochondrial respiratory capacity and is required for cold-induced activation of the thermogenic gene program in BAT. Mitochondrial fatty acid β-oxidation (FAO) is critical for maintaining the brown adipocyte phenotype both during times of activation and quiescence. FAO also fuels the increase in uncoupled mitochondrial respiration and contributes to inducing the expression of thermogenic genes such as *Ucp1, Ppargc1α*, and *Dio2* in response to adrenergic stimulation (26, 27). Consequently, mice with adipose-specific deficits in FAO are severely cold-intolerant, demonstrating its role in cold-induced thermogenesis (13, 14).

Although, not directly tested, it is likely that reduced FAO in OPA1 BAT KO mice (KO) contributed to the impaired thermogenic activation of BAT. However, in contrast to many models of mitochondrial dysfunction or FAO defects in BAT, KO mice displayed improved cold adaptation. The increased core body temperatures at thermoneutrality and heightened tolerance to cold at 4°C correlated with a significant increase in compensatory browning of subcutaneous white adipose tissue (WAT) in KO mice. Several studies reported increased browning of WAT following BAT impairment. Surgical removal of inter-scapular BAT (iBAT) enhanced WAT browning due to increased sympathetic input to WAT in mice, which was accompanied by reduced adiposity (28). In addition, denervation of iBAT increased sympathetic input to subcutaneous fat to induce compensatory browning (18). Genetic models of BAT paucity demonstrated a similar phenomenon. Ablation of type 1A BMP-receptor (*Bmpr1A*) in brown adipogenic progenitor cells induces a severe paucity of BAT, which increased sympathetic input to WAT, thereby promoting browning, and maintaining normal temperature homeostasis and resistance to DIO (18). Lastly, ablating insulin receptor in the Myf5 lineage reduced interscapular BAT mass in mice that maintained normal thermogenesis on the basis of compensatory browning of sub-cutaneous WAT and increased lipolytic activity in BAT (17). We, therefore postulated that OPA1 deletion in BAT increased sympathetic input to WAT to promote compensatory browning. Indeed, we observed increased tyrosine hydroxylase protein levels in WAT, even at room temperature conditions, suggesting increased sympathetic innervation of sub-cutaneous fat. This compensatory browning likely contributes to increased core body temperature at both 30°C and at 4°C.

The mechanisms linking browning of WAT when BAT function is impaired are incompletely understood. Our data in KO mice demonstrated that FGF21 secretion from BAT, is independent of BAT thermogenic activation, and mediates this adaptation. FGF21 release from activated BAT might contribute to many of the metabolic benefits associated with BAT activity (29). Furthermore, FGF21 is a key regulator of WAT browning in mice, leading to increased thermogenesis and energy expenditure (19). Our data in mice lacking OPA1 and FGF21 in BAT (OPA1/FGF21 DKO) revealed that BAT-derived FGF21 is required for increased browning of WAT in KO mice under isocaloric conditions. These data support the idea that FGF21 is a potent inducer of WAT browning, and demonstrates that BAT-derived FGF21 mediates compensatory browning of WAT following OPA1 deletion in BAT, that contributes to the metabolic phenotype of these mice, despite absence of BAT activation.

Pharmacological administration of FGF21 increases energy expenditure and thermogenic gene expression in BAT and WAT (19, 30). However, the role of endogenous BAT-derived FGF21 upon cold exposure remains incompletely understood. In KO mice, BAT-derived FGF21 was required for the baseline increase in core body temperature and for the resistance to cold stress. This result contrasts with a recent report demonstrating that FGF21 plays a negligible role in the systemic adaptations to long-term cold exposure in mice, including browning of subcutaneous WAT, in a global FGF21 knock out model (31). Conversely, Fisher et al. showed that whole body ablation of FGF21 impaired the response to cold stress, when placing mice from 27°C to 5°C for 3 days (19). These studies utilized global knock out models, thus, the source of FGF21 is unclear. Moreover, liver-derived, but not adipose tissue-derived FGF21, was shown to enter the circulation within the first hours of cold exposure, contributing to thermoregulation, via its action in the central nervous system (32). Together, these studies suggest that BAT-derived FGF21 could be dispensable for thermoregulation during short- and long-term cold acclimation in wild type mice. However, our data clearly demonstrates that BAT-derived FGF21 does mediate the compensatory response of mice lacking OPA1 in BAT after short-term cold exposure, as OPA1/FGF21 DKO mice undergo a steep decline in core body temperature when exposed to cold. Our model results from constitutive deletion of the *Opa1* gene. Therefore, our phenotype could reflect the effects of long-term exposure to persistent mildly increased endogenous FGF21 circulating levels, leading to considerable browning of WAT even at ambient temperature conditions. This augmented thermogenic response in WAT could prime these animals to better adapt to cold temperatures and might also contribute to the increased resting metabolic rates and leanness in KO mice.

The mechanisms governing FGF21-mediated browning are incompletely understood. FGF21 may directly promote browning of WAT in part, by induction of *Ppargc-1α* (19), and contributes to the induction of WAT browning during cold acclimation (28). However, central or peripherally administered FGF21 failed to induce beige fat in mice lacking β-adrenoceptors, indicating the requirement for an intact adrenergic system (33, 34). Lastly, liver-derived FGF21 mediates thermoregulation via its action in the central nervous system, rather than in adipose tissue (32). Together, these data suggest that FGF21 might signal centrally to activate the sympathetic nervous system and promote browning in KO mice. However, we observed increased tyrosine hydroxylase levels in WAT of OPA1/FGF21 DKO mice, which lacked the compensatory browning of WAT. This indicates that BAT-derived FGF21 does not mediate the increase in sympathetic innervation of WAT in KO mice, and that increased sympathetic input to WAT is not sufficient to promote browning in the absence of elevated circulating FGF21 levels in mice. Thus, FGF21-independent mechanisms may mediate the compensatory increase in sympathetic tone to WAT in KO mice. However, FGF21 and the sympathetic nervous system may act cooperatively to induce browning in KO mice fed isocaloric diet. Finally, tyrosine hydroxylase levels were reduced in iWAT of both KO and OPA1/FGF21 DKO mice fed HFD, while UCP1 levels were significantly elevated, suggesting that increased sympathetic innervation is dispensable for WAT browning and for the resistance to DIO in KO and DKO mice fed HFD.

FGF21 is strongly induced in BAT in response to thermogenic stimulation, via mechanisms that involve β-adrenergic signaling activation (19). Because BAT thermogenic function was impaired in KO mice, and FGF21 levels were elevated even at ambient temperatures conditions, we hypothesized that alternative signaling pathways regulated FGF21 induction in KO mice. We and others have shown that OPA1 deletion in muscle leads to FGF21 induction via activation of the ER stress pathway (35, 36). ATF4, a transcription factor downstream of the unfolded protein response (UPR), has been proposed to induce transcriptional regulation of FGF21 in models of mitochondrial stress, as part of the integrated stress response (37). Our data in mice lacking both OPA1 and ATF4 selectively in BAT demonstrated that ATF4 is required for FGF21 induction in and secretion from BAT in response to OPA1 deletion. Accordingly, lack of ATF4 in BAT recapitulated the effects of FGF21 deficiency in OPA1 BAT KO mice, including lack of baseline browning, normalized metabolic rates and impaired adaptive thermogenesis. Taken together, our data strongly suggests that ATF4, likely downstream of ER stress activation, is required for FGF21 induction in OPA1 BAT KO mice. Interestingly, a recent study demonstrated induction of ER stress and ATF4 in BAT in response to cold stress (38). Future studies focusing on the role of ATF4 in BAT thermogenesis and BATokine secretion might identify mechanisms for BAT-mediated metabolic adaptations that might be independent of β-adrenergic stimulation of BAT.

Surprisingly, BAT-derived FGF21 does not appear to mediate the resistance to DIO and insulin resistance (IR) in KO mice. These data suggest that factors other than FGF21 may contribute to the lean phenotype in these mice, may and mediate the increase in metabolic rates and browning of WAT when mice are fed HFD. Indeed, the diet-induced increase in FGF21 circulating levels was completely blunted in OPA1 BAT KO mice. This finding further supports the conclusion that FGF21 is not required for the systemic adaptations observed when OPA1 BAT KO mice are fed an obesogenic diet. Future studies investigating BAT secretome in this mouse model might identify additional BAT-derived secreted factors, which could potentially mediate the resistance to DIO. It is also possible that alternative mechanisms, independent of BAT secretome, participate in mediating this metabolic protection in OPA1 BAT KO mice.

In conclusion, these studies reveal an important stress response pathway in BAT in the absence of OPA1, consisting of induction of FGF21 as a BATokine via ATF4-dependent mechanisms, promoting leanness, and improving thermoregulation when BAT function is dampened. Determining the physiological relevance of this ATF4-FGF21 axis in BAT physiology and BAT-mediated metabolic adaptations may lead to novel therapeutic approaches to combat obesity and associated disorders.

## Material and Methods

### Mouse Models

Experiments were performed in male and/or female mice on a C57Bl/6J background. OPA1^fl/fl^ mice (39), FGF21^fl/fl^ mice (40), and ATF4^fl/fl^ mice were (41) were generated as previously described. Transgenic mice expressing cre recombinase under the control of the *Ucp1* promoter (Tg (Ucp1-cre)1Evdr) (16), and transgenic mice expressing a tamoxifen-inducible cre under the control of the *Adipoq* gene promoter (C57BL/6-Tg(Adipoq-cre/ERT2)1Soff/J) (42) were acquired from the Jackson Laboratories (#024670 and #025124, respectively). Mice were weaned at 3 weeks of age and were either kept on standard chow (2029X Harlan Teklad, Indianapolis, IN, USA) or were fed special diets. For diet-induced obesity studies, 6-week old mice were divided into a control-diet group (Cont; 10% Kcal from fat—Research Diets, New Brunswick, NJ, USA, D12450J) or a high-fat diet group (HFD; 60% Kcal from fat—Research Diets D12492) and were kept on these respective diets for 12 weeks. For the cold exposure experiments, mice were acclimated to 30°C (thermoneutral temperature for mice) for 7 days prior to being cold-exposed. Animals were housed at 22°C with a 12-h light, 12-h dark cycle with free access to water and standard chow or special diets, unless otherwise noticed. All mouse experiments presented in this study were conducted in accordance with the animal research guidelines from NIH and were approved by the University of Iowa IACUC.

## Methods Details

### Studies with mice reared at thermoneutrality

A subset of OPA1 BAT KO female mice and their wild type (WT) littermate controls were transferred to a rodent environmental chamber (Power Scientific) set at 30 °C following weaning (∼ 4 weeks of age), and were housed for the subsequent 4 weeks (until ∼ 8 weeks of age). Body composition and indirect calorimetry were measured at the end of the 4 weeks. Mice were transferred to an OxyMax Comprehensive Lab Animal Monitoring System (CLAMS, Columbus Instruments International), where oxygen consumption, food intake and ambulatory activity were measured. Body composition was determined by NMR, and brown adipose tissue (BAT), inguinal white adipose tissue (iWAT) and gonadal white adipose tissue (gWAT) depots were weighed upon tissue harvest.

### Cold exposure experiments

Core body temperature telemeters (Respironics, G2 E-Mitter, Murrysville, PA, USA) were surgically implanted into the abdominal cavity of 8-10-week old male mice, and mice were then allowed to recover for 6 days post-surgery, while individually housed in a rodent environmental chamber (Power Scientific) at 30 °C. Mice were then transferred to an OxyMax Comprehensive Lab Animal Monitoring System (CLAMS, Columbus Instruments International) at 30 °C for 4 days, followed by 4 °C for 3 days, as previously described (19). Core body temperature was recorded every 17 minutes throughout the experiment, along with O_2_ and CO_2_ levels, food intake and ambulatory activity, as estimated by photoelectric beam breaks in the X + Y plane. For the acute cold exposure experiments, 8-week old mice were initially individually housed in the rodent environmental chamber at 30 °C for 7 days. The initial temperature (t0) was recorded using a rectal probe (Fisher Scientific, Lenexa, KS, USA) at 7 am of day 8, after which the temperature was switched to 4 °C. Once the desired temperature was reached, we recorded rectal temperatures hourly for up to 4 hours of cold exposure.

### Glucose and insulin tolerance tests, nuclear magnetic resonance, and serum analysis

Glucose tolerance tests (GTT) were performed after a 6-h fast, and mice were administered glucose intraperitoneally (2 g/kg body weight), as described (43). Insulin tolerance tests (ITT) were performed after a 2-h fast by injecting insulin intraperitoneally (0.75 U/kg body weight; Humulin, Eli Lilly, Indianapolis, IN, USA). Blood glucose was determined using a glucometer at regular time intervals (Glucometer Elite; Bayer, Tarrytown, NY, USA). Insulin solution was prepared in sterile 0.9% saline and dosages were based on body weight. Plasma insulin was measured after a 6-hour fast using a commercially available kit according to the manufacturers’ directions (Ultra-Sensitive Mouse Insulin ELISA Kit, Chrystal Chem, Downers Grove, IL, USA). Serum FGF21 (BioVendor ELISA kit, Asheville, NC, USA) were measured using commercially available kits according to the manufacturers’ directions. Whole body composition was measured by nuclear magnetic resonance in the Bruker Minispec NF-50 instrument (Bruker, Billerica, MA, USA). NMR was performed at the University of Iowa Fraternal Order of Eagles Diabetes Research Center Metabolic Phenotyping Core.

### Analysis of triglyceride levels

Triglycerides levels were measured in liver and in serum collected after a 6-h fast using the EnzyChrom™ Triglyceride Assay Kit (BioAssay Systems, Hayward, CA, USA). Liver triglycerides were extracted using a solution of isopropanol and Triton X-100, as recommended by the manufacturers (44).

### RNA extraction and quantitative RT–PCR

Total RNA was extracted from tissues with TRIzol reagent (Invitrogen) and purified with the RNeasy kit (Qiagen Inc, Germantown, MD, USA). RNA concentration was determined by measuring the absorbance at 260 and 280 nm using a spectrophotometer (NanoDrop 1000, NanoDrop products, Wilmington, DE, USA). Total RNA (1 μg) was reverse-transcribed using the High-Capacity cDNA Reverse Transcription Kit (Applied Biosystems, Waltham, MA, USA), followed by qPCR reactions using SYBR Green (Life Technologies, Carlsbad, CA, USA) (35). Samples were loaded in a 384-well plate in triplicate, and real-time polymerase chain reaction was performed with an ABI Prism 7900HT instrument (Applied Biosystems, Waltham, MA, USA). The following cycle profile was used: 1 cycle at 95°C for 10 min; 40 cycles of 95°C for 15 s; 59°C for 15 s, 72°C for 30 s, and 78°C for 10 s; 1 cycle of 95°C for 15 s; 1 cycle of 60°C for 15 s; and 1 cycle of 95°C for 15 s. Data were normalized to *Gapdh* expression and results are shown as relative mRNA levels. qPCR primers were designed using Primer-Blast or previously published sequences (45). Utilized primers are listed in Table 1.

**Table 1.**
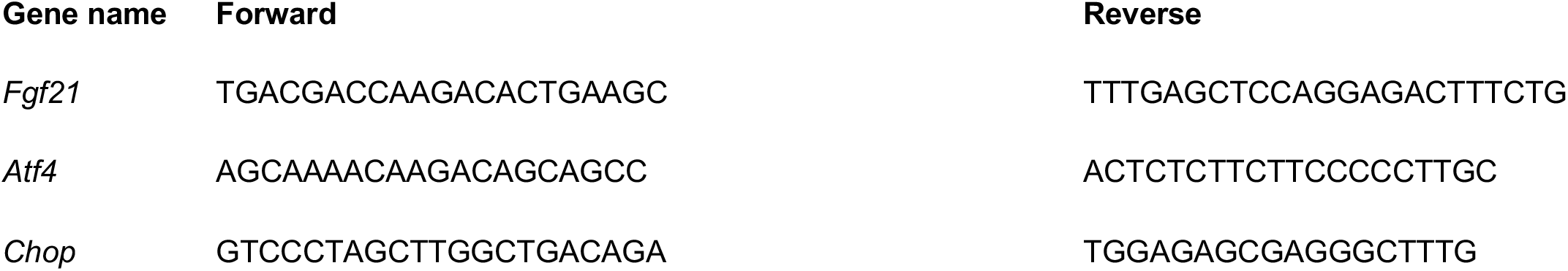
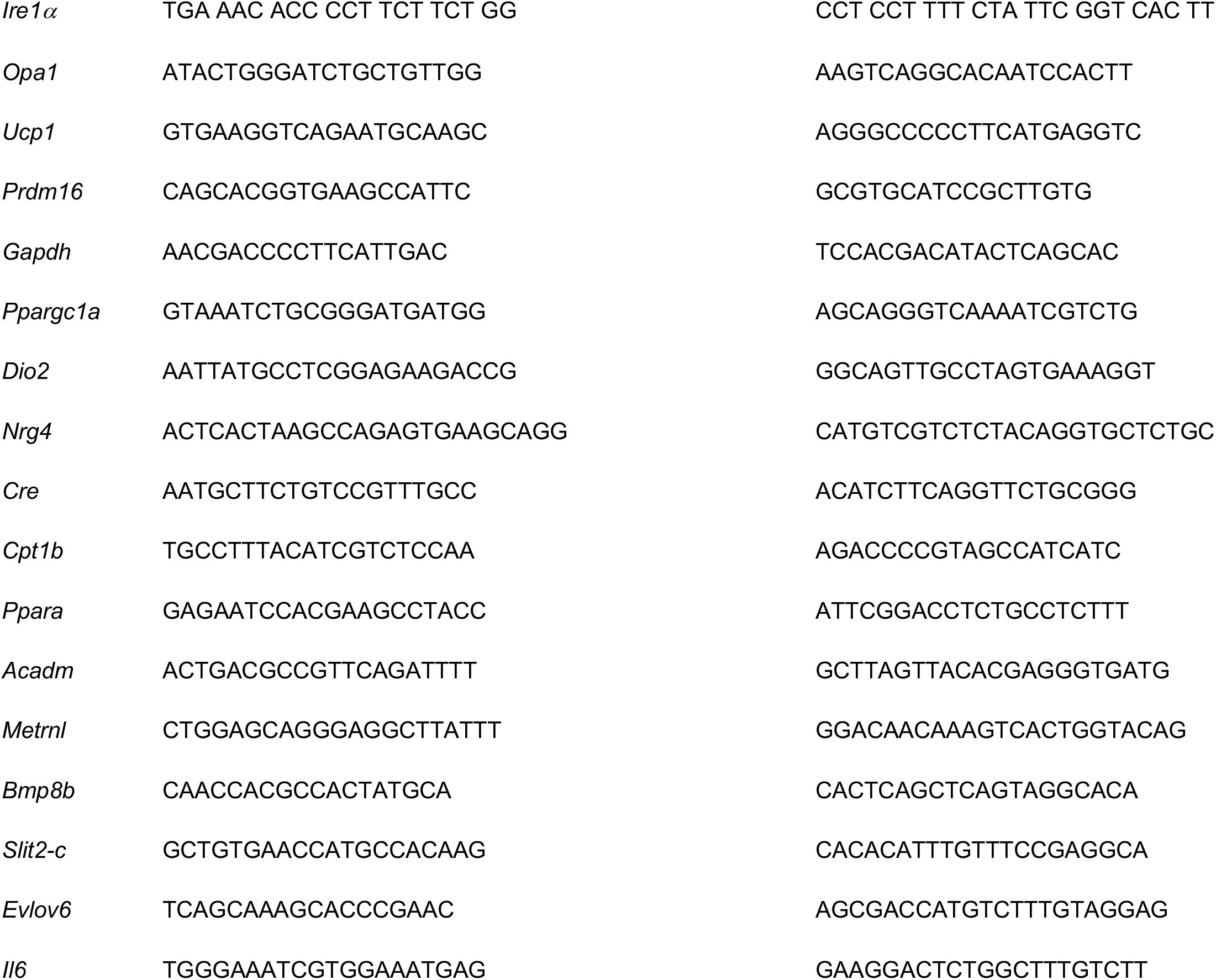
Primers

### Western blot analysis

Immunoblotting analysis was performed as previously described (46). Approximately, 50 mg of frozen tissue was homogenized in 200 μl lysis buffer containing (in mmol/l) 50 HEPES, 150 NaCl, 10% glycerol, 1% Triton X-100, 1.5 MgCl_2_, 1 EGTA, 10 sodium pyrophosphate, 100 sodium fluoride, and 100 μmol/l sodium vanadate. Right before use, HALT protease/phosphatase inhibitors (Thermo Fisher Scientific, Waltham, MA, USA) were added to the lysis buffer and samples were processed using the TissueLyser II (Qiagen Inc., Germantown, MD, USA). Tissue lysates or freshly isolated mitochondria were resolved on SDS–PAGE and transferred to nitrocellulose membranes (Millipore Corp., Billerica, MA, USA). Membranes were incubated with primary antibodies overnight and with secondary antibodies for 1 h, at room temperature.

## Antibodies

Primary Antibodies: OPA1 (1:1,000, BD Biosciences, San Jose, CA, USA, #612606), FGF21 (1:1,000, Abcam, Cambridge, UK, #ab171941), VDAC (1:1,000, Thermo Scientific, #PA1-954A), GAPDH (1:1,000, Cell Signaling Technology, Danvers, MA, USA, #2118), UCP1 (1:1,000, Abcam, #Ab10983), SDH (1:1,000, Abcam, #Ab14714), *α*-tubulin (1:1,000, Sigma, St. Lois, MO, #T9026), *β*-actin (1:1,000,Sigma, #A2066), tyrosine hydroxylase (1:1,000, Cell Signaling Technology, #2792), phosphorylated eIF2*α* serine 51 (1:1,000, Cell Signaling Technology, #3597) and eIF2α (1:1,000, Santa Cruz Biotechnology, Dallas, TX, USA, #SC81261). Secondary antibodies: IRDye 800CW anti-mouse (1:10,000, LI-COR, Lincoln, NE, USA, #925-32212) and Alexa Fluor anti-rabbit 680 (1:10,000, Invitrogen #A27042). Fluorescence was quantified using the LiCor Odyssey imager.

## Mitochondrial Isolation

Mitochondrial fraction was isolated from interscapular brown adipose tissue (iBAT) or from inguinal white adipose tissue (iWAT), as previously described (47). Briefly, tissue was excised, rinsed in ice-cold PBS and maintained in ice-cold isolation buffer (500 mM EDTA, 215 mM D-mannitol, 75 mM sucrose, 0.1% free-fatty acid bovine serum albumin, 20 mM HEPES, pH 7.4 with KOH) until ready for homogenization. Bradford assay was performed to determine the protein concentration.

### Oxygen consumption

Mitochondrial function was assessed using Seahorse XF96 analyzer. Briefly, 2.5ug of BAT or iWAT mitochondria were seeded on a Polyethylene Terephthalate (PET) plate and centrifuged for 20 minutes at 2000xg at 4°C. Substrates were added to the assay buffer at the following final concentrations-Pyruvate at 2mM, Malate at 0.8mM and Palmitoyl carnitine at 0.02mM. Measurements were performed at baseline conditions (State 2) or after ADP (2mM) injection (State 3). Substrates were freshly prepared and reagents were purchased from Sigma (St. Louis, MO, USA).

### ATP synthesis rates

ATP synthesis rates were assessed in 20 µg of mitochondria isolated from iBAT using a fluorometer (Horiba Systems, Irvine, CA, USA). Briefly, buffer Z lite (105mM KMES, 30mM KCl, 10mM KH_2_PO_4_, 5mM MgCl_2_.6H_2_O, 0.5mg/mL BSA, pH 7.4 with KOH) supplemented with glucose, hexokinase/G-6-PDH (Sigma, St. Louis, MO, USA), Ap5a (an inhibitor of adenylate kinase, Sigma, St. Louis, MO, USA) and NADP (Sigma, St. Louis, MO, USA) was added with 20 µg of mitochondria. ADP (75 µM) stimulated ATP synthesis rates were measured kinetically in the presence of palmitoyl carnitine (5 µM) and malate (1.6 mM) from the formation of NADPH in a coupled reaction that uses ATP for conversion of glucose to glucose-6-phosphate (G6P) by hexokinase and subsequently to 6-phosphogluconolactone by G6P dehydrogenase coupled with reduction of NADP to NADPH. NADPH accumulation is measured by fluorometry using excitation/emission wavelengths of 345nm/460nm respectively (48).

## Transmission Electron Microscopy

Electron micrographs of iBAT and iWAT were prepared as previously described (35). Briefly, iBAT and iWAT were trimmed into tiny pieces using a new blade to minimize mechanical trauma to the tissue. Tissues were fixed overnight (in 2% formaldehyde and 2.5% glutaraldehyde) rinsed (0.1% cacodylate pH 7.2) and stained with increasing concentrations of osmium (1.5%, 4%, 6%), and dehydrated with increasing concentrations of acetone (50%, 75%, 95%, 100%). Samples were then embedded, cured, sectioned and poststained with uranyl and lead. Sections were then imaged on Jeol 1230 Transmission electron microscope.

## Histology and Immunohistochemistry

Fragments of BAT and iWAT were embedded in paraffin, portioned into 5-μm-thick sections, and stained with hematoxylin-eosin (Fisher, Pittsburgh, PA, USA). For immunohistochemistry, iWAT sections were deparaffinized, re-hydrated, blocked with 10% goat serum and incubated overnight with a rabbit primary antibody against UCP1 (Abcam, Ab1098, 1:250). Sections were, then, incubated with an anti-rabbit biotinylated secondary antibody (1:500) for 1 hour, incubated with peroxidase streptavidin solution (1:500) for 30 min and revealed with DAB chromogen solution for 10 seconds (Vector Laboratories, Burlingame, CA, USA). Light microscopy was performed using a Nikon Eclipse Ti-S microscope (Nikon, Melville, NY, USA).

## Cell culture and treatments

Brown adipose tissue stromal vasculature fraction (SVF) was isolated from 6-day old OPA1 floxed mice harboring the tamoxifen-inducible cre recombinase, Cre-ERT2, under the control of the *Adipoq* gene promoter. Cells were grown and differentiated as previously described (49). After cells were fully differentiated, brown adipocytes were treated with 500 nM 4-hydroxytamoxifen (Sigma, St. Louis, MO, USA) for 72 hours to induce OPA1 deletion (KO cells) or with vehicle solution (WT cells). Cells were then switched to serum-free and phenol-free DMEM/F12 for 6 hours, after which the media was collected for FGF21 measurements. Cells were washed with ice-cold PBS and harvested for subsequent analysis.

### Data analysis

Unless otherwise noted, all data are reported as mean ± SEM. Student’s *t*-test was performed for comparison of two groups, and Two-Way ANOVA followed by Tukey multiple comparison test was utilized when more than three groups were compared. A probability value of *P* ≤ 0.05 was considered significantly different. Statistical calculations were performed using the GraphPad Prism software (La Jolla, CA, USA).

## Acknowledgment

This work was supported by grants HL127764 and HL112413 from the NIH, 20SFRN35120123 from the American Heart Association (AHA) and the Teresa Benoit Diabetes research fund to E.D.A., who is an established investigator of the AHA; by AHA Scientist Development Grant 15SDG25710438 and NIH DK125405 to R.O.P.; by the Diabetes Research Training Program funded by the NIH (T32DK112751-01) to S.H.B; and by the NIH 1R25GM116686 to L.M.G.P. Metabolic phenotyping was performed at the Metabolic Phenotyping Core at the Fraternal Order of Eagles Diabetes Research Center. Analysis of mRNA expression was performed by qPCR at the Genomics Division at The Iowa Institute of Human Genetics. Electron Microscopy was performed at the Central Microscopy Research Facility at the University of Iowa. We would like to thank Dr. Hiromi Sesaki, at John Hopkins, for providing us with the OPA1 floxed mice.

## Author Contributions

R.O.P. and E.D.A conceived the project and coordinated all aspects of this work. R.O.P. conceptualized, designed and conducted the experiments, analyzed data and wrote the manuscript. A.M., A.C.O, S.H.B, S.M.T, E.T.W. and M.W. conducted the experiments and analyzed data. P. P., A.D.T., R.H., W.B.T., L.M.G.P aided with animal work and processed tissues for biochemical analysis. M.M. performed histology, and R.A.S. performed TEM; C.M.A. and M.J.P. provided essential materials and critical expertise. E.D.A edited the manuscript.

## Supplemental Information

**Figure S1:**
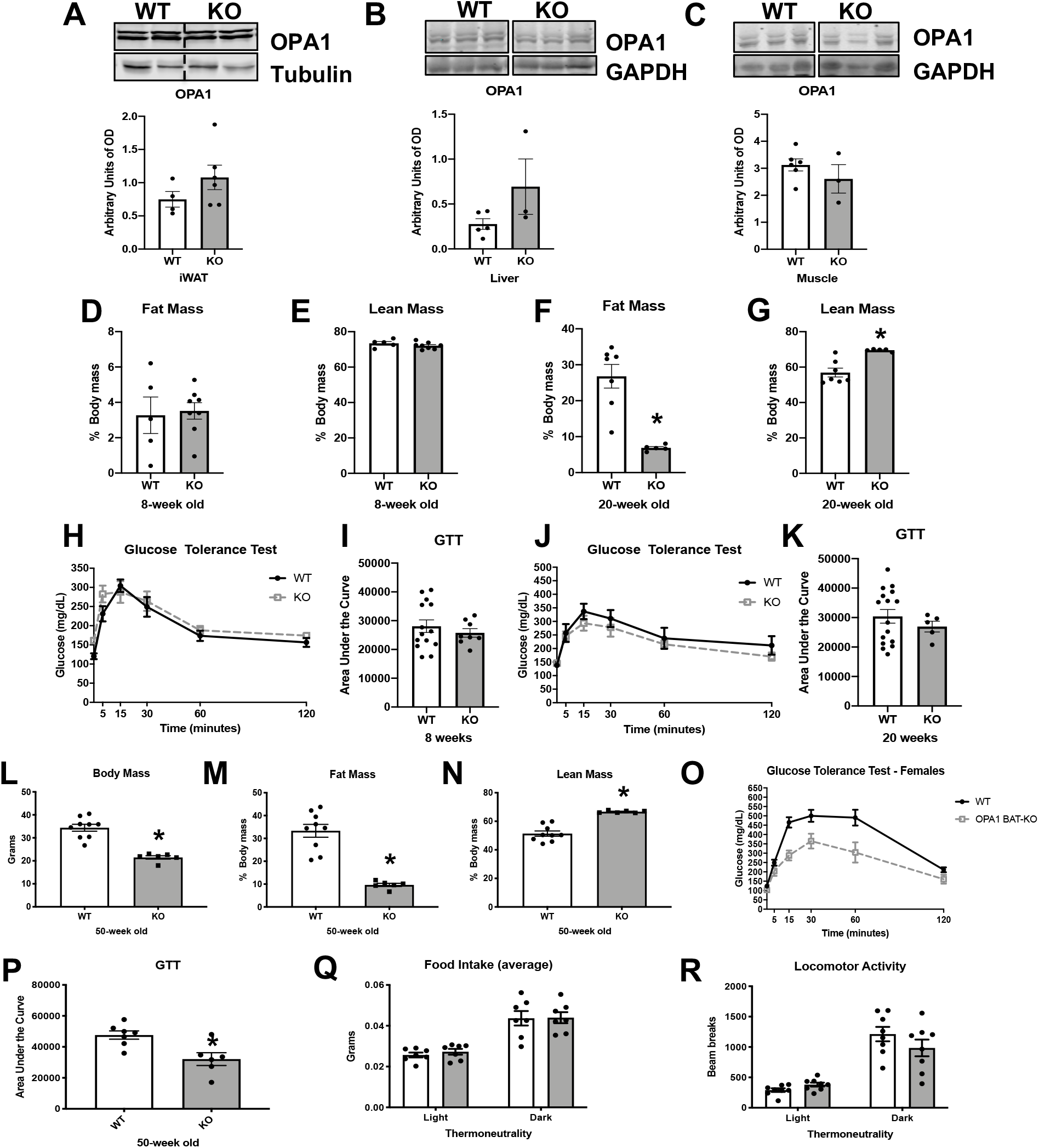
Age-dependent changes in body composition and glucose homeostasis in OPA1 BAT KO mice. Related to figure 1. **A-C**. Representative immunoblot of OPA1 and densitometric analysis of OPA1 normalized by tubulin or GAPDH. **A**-Inguinal White adipose tissue (iWAT) (dashed line separates genotypes). **B**-Liver (images were cropped from the same membrane). **C**-Skeletal muscle. **D-K**. Body composition in 8 and 20-week-old mice. **D**-Percent fat mass to body weight at 8 weeks. **E**-Percent lean mass to body weight at 8 weeks. **F**. Percent fat mass to body weight at 20 weeks. **G**. Percent lean mass to body weight at 20 weeks. **H**. Glucose tolerance test (GTT) in 8-week-old mice. **I**. Area under the curve for the GTT performed at 8 weeks. **J**. GTT in 20-week-old mice. **K**. Area under the curve for the GTT performed at 20 weeks. **L-P**. Body composition and glucose homeostasis in 50-week-old female mice. **L**. Body mass. **M**. Percent fat mass to body weight. **N**. Percent lean mass to body weight. **O**. GTT in 50-week-old mice. **P**. Area under the curve for the GTT performed at 50 weeks. **Q**. Food intake measured in 8-week-old mice under thermoneutral conditions. **R**. Locomotor activity measured in 8-week-old mice under thermoneutral conditions. Data are expressed as means ± SEM. Data are expressed as means ± SEM. Significant differences were determined by Student’s *t*-test, using a significance level of *P* < 0.05. (*) Significantly different vs. WT mice.

**Figure S2:**
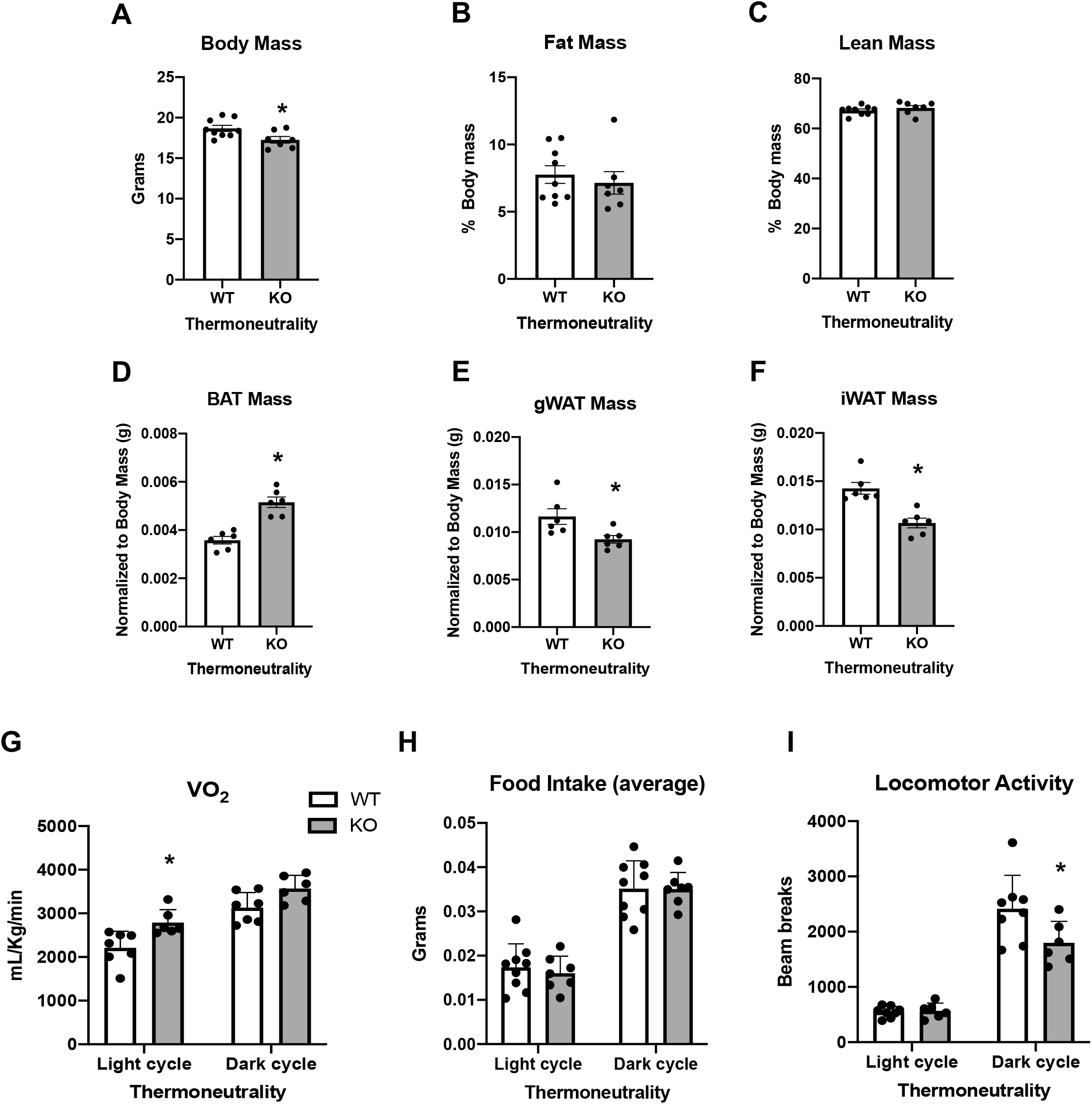
Data collected in 8-week-old OPA1 BAT KO mice (KO) and their wild type littermate controls (WT) reared at thermoneutrality. Related to figure 1. **A**. Body mass. **B**. Percent fat mass normalized to body weight. **C**. Percent lean mass normalized to body weight. **D**. BAT mass normalized by body weight. **E**. gWAT mass normalized by body weight. **F**. iWAT mass normalized to body weight. **G**. Oxygen consumption. **H**. Average food intake. **I**. Locomotor activity. Data are expressed as means ± SEM. Significant differences were determined by Student’s *t*-test using a significance level of *P* < 0.05. (*) Significantly different vs. WT mice.

**Figure S3:**
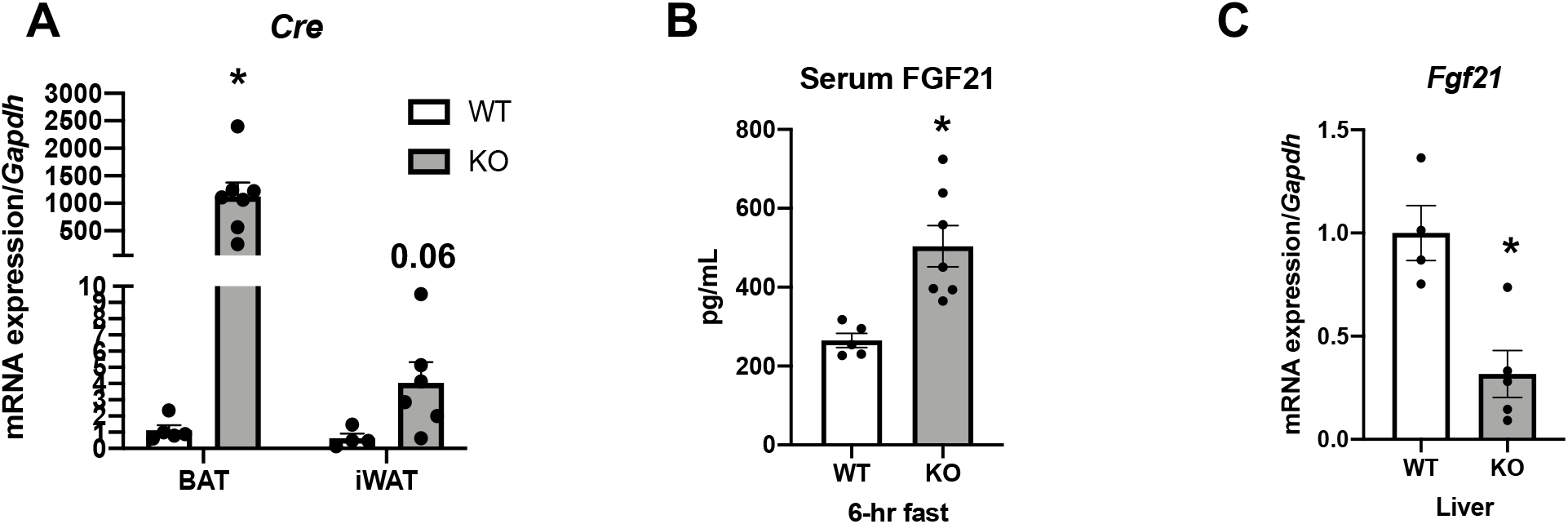
Data collected in 8-week-old OPA1 BAT KO mice (KO) and their wild type littermate controls (WT) at room temperature conditions. Related to figure 3. **A**. *Cre* mRNA expression in BAT and iWAT normalized to *Gapdh* expression. **B**. Fasting FGF21 serum levels **C**. *Fgf21* mRNA expression in livers normalized to *Gapdh* expression. Data are expressed as means ± SEM. Significant differences were determined by Student’s *t*-test using a significance level of *P* < 0.05. (*) Significantly different vs. WT mice.

**Figure S4:**
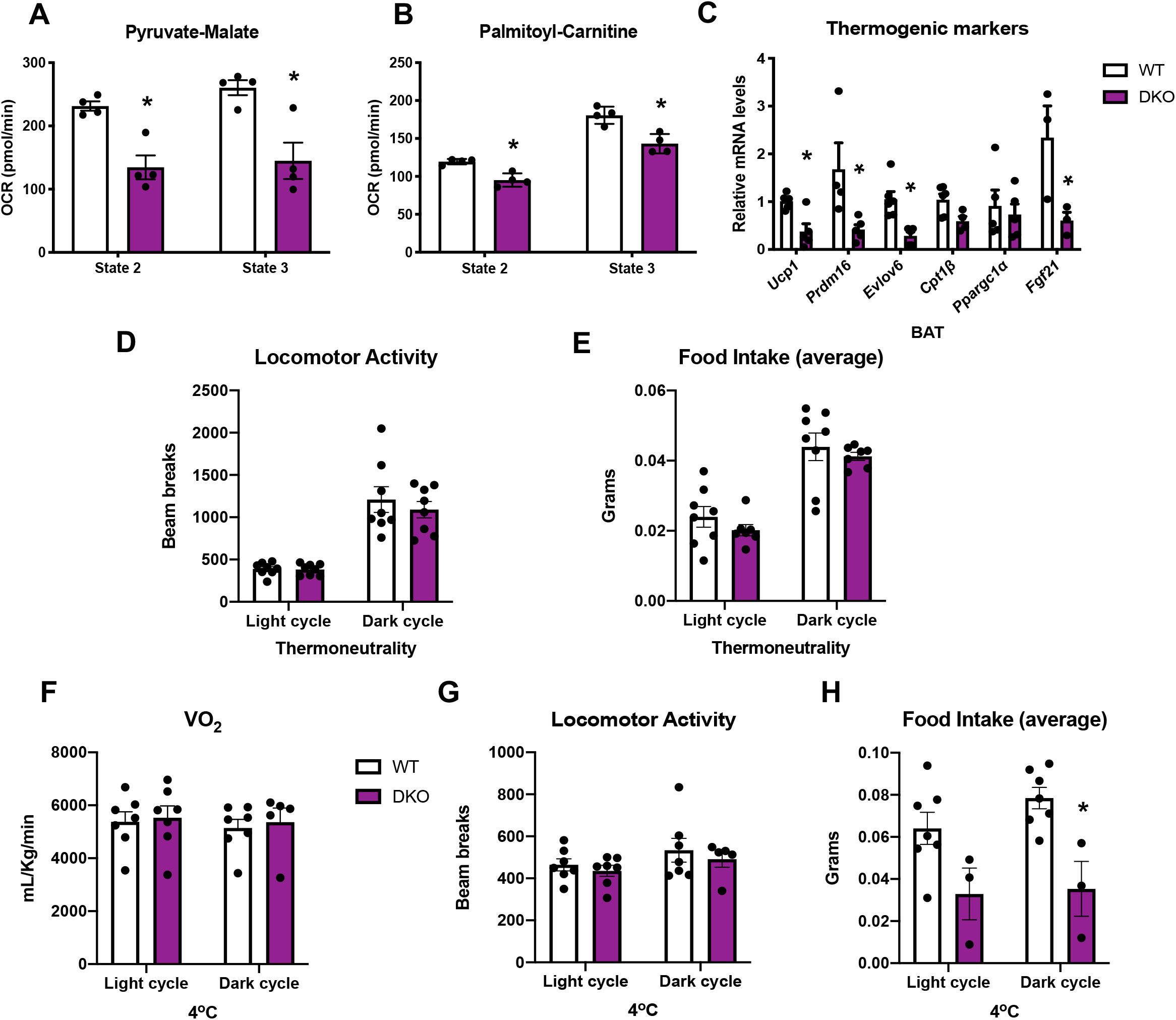
Data collected in OPA1/FGF21 BAT DKO mice and their wild type littermate controls (WT). Related to figure 4. **A**. State 2 and state 3 pyruvate-malate-supported oxygen consumption rates (OCR) in mitochondria isolated from BAT. **B**. State 2 and state 3 palmitoyl-carnitine-supported OCR in mitochondria isolated from BAT. **C**. mRNA expression of thermogenic genes in BAT tissue collected from mice raised at room temperature (22oC). **D**. Locomotor activity in 10-week-old male mice measured at thermoneutrality (30oC). **E**. Average food intake in 10-week-old male mice measured at thermoneutrality. **F**. Oxygen consumption measured in 10-week-old male mice housed at 4oC for 3 days. **G**. Locomotor activity measured in 10-week-old male mice housed at 4oC for 3 days. **H**. Average food intake measured in 10-week-old male mice housed at 4oC for 3 days. Data are expressed as means ± SEM. Significant differences were determined by Student’s *t*-test, using a significance level of *P* < 0.05. (*) Significantly different vs. WT mice.

**Figure S5:**
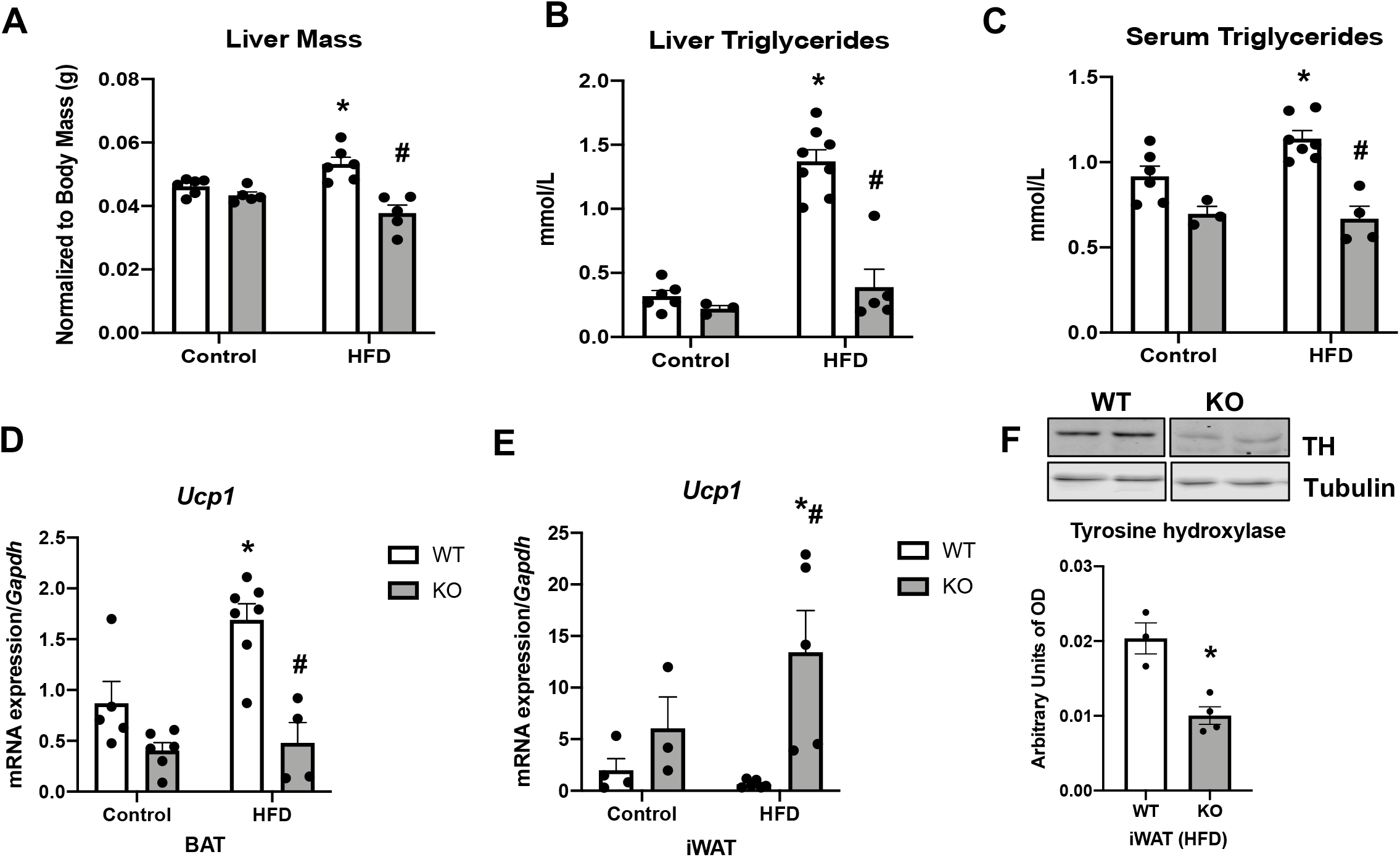
Data collected in OPA1 BAT KO mice (KO) and their wild type littermate controls (WT) fed either control (10% fat content) or HFD (60% fat content) for 12 weeks. Related to figure 5. **A**. Liver mass. **B**. Liver triglycerides levels. **C**. Serum triglycerides levels. **D**. mRNA expression of *Ucp1* in BAT. **E**. mRNA expression of *Ucp1* in iWAT. **F**. Representative immunoblot of tyrosine hydroxylase (TH) levels in iWAT of mice fed a HFD and densitometric quantification normalized by tubulin (images were cropped from the same membrane). Data are expressed as means ± SEM. Significant differences were determined by Two-Way ANOVA, using a significance level of *P* < 0.05. (*) Significantly different vs. WT control, (#) significantly different vs. WT HFD.

**Figure S6:**
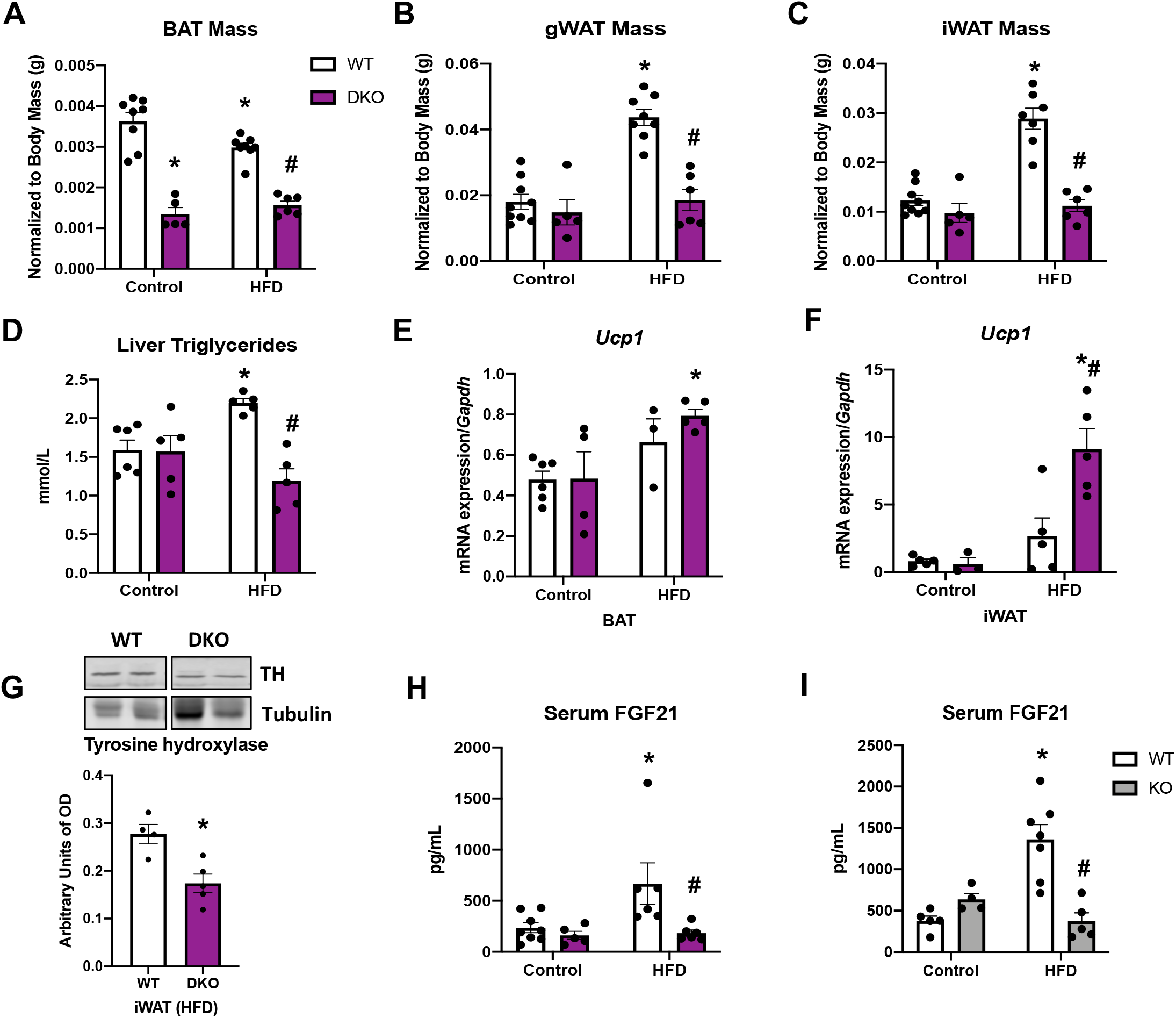
Data collected in OPA1 BAT KO mice (KO), OPA1/FGF21 DKO mice or their respective wild type littermate controls (WT) fed either control (10% fat content) or HFD (60% fat content) for 12 weeks. Related to figure 6. **A**. BAT mass normalized to body mass. **B**. gWAT mass normalized to body mass. **C**. iWAT mass normalized to body mass. **D**. Liver triglyceride levels. **E**. mRNA expression of *Ucp1* in BAT normalized to *Gapdh*. **F**. mRNA expression of *Ucp1* in iWAT. **G**. Representative immunoblot of tyrosine hydroxylase (TH) levels in iWAT of mice fed a HFD and densitometric quantification normalized by tubulin (images were cropped from the same membrane). **H**. Serum FGF21 level in WT and DKO mice. **I**. Serum FGF21 levels in WT and OPA1 BAT KO mice (KO). Data are expressed as means ± SEM. Significant differences were determined by Two-Way ANOVA, using a significance level of *P* < 0.05. (*) Significantly different vs. WT control, (#) significantly different vs. WT HFD.

